# Integrated RNA-seq analysis identifies ABC transporters mediating taxane export in *Taxus* species

**DOI:** 10.64898/2026.05.10.723993

**Authors:** Jaber Nasiri, Alireza Fotuhi Siahpirani, Yueming Dong, Catherine Xu, Yu Xia, Codruta Ignea

**Author notes:** Contributed equally to this work (Jaber Nasiri, Alireza Fotuhi Siahpirani). Corresponding author: Codruta Ignea.

## Abstract

RNA-seq datasets from medicinal yews are crucial for studying paclitaxel biosynthesis. However, cross-study data analyses are hindered by pronounced batch effects. Here, we compiled 45 RNA-seq samples from three studies across four tissues (bark, leaf, root, stem) and assessed 35 preprocessing pipelines combining six normalization strategies with five batch-effect correction approaches. Unsupervised clustering (HCA, k-means, Grade-of-Membership), evaluated using Jaccard and Adjusted Rand indices, revealed significant variability in batch effect removal. Supervised classification of tissue and project labels (Random Forest and linear/radial SVM) demonstrated improved accuracy in tissue type prediction, highlighting the effectiveness of correction methods. The processed data facilitated the identification of 189 putative ABC transporters across samples, six of which showing a strong correlation to the gene encoding 10-deacetylbaccatin-III-10β-O-acetyltransferase, a key biosynthetic enzyme in the taxol pathway. High expression levels in leaf and bark further support their role in taxane intermediates trafficking in taxol biosynthesis. Structural analysis and molecular docking further supported the selection of these candidates, and the agreement between transcriptomic ranking and docking-based prioritization suggests that these transporters may participate in taxane intermediate recognition, trafficking, or export. These findings demonstrate the importance of normalization and batch effect correction in RNA-seq analysis to advance gene discovery in *Taxus* species and, more broadly, in plant research.

**Graphical abstract:** 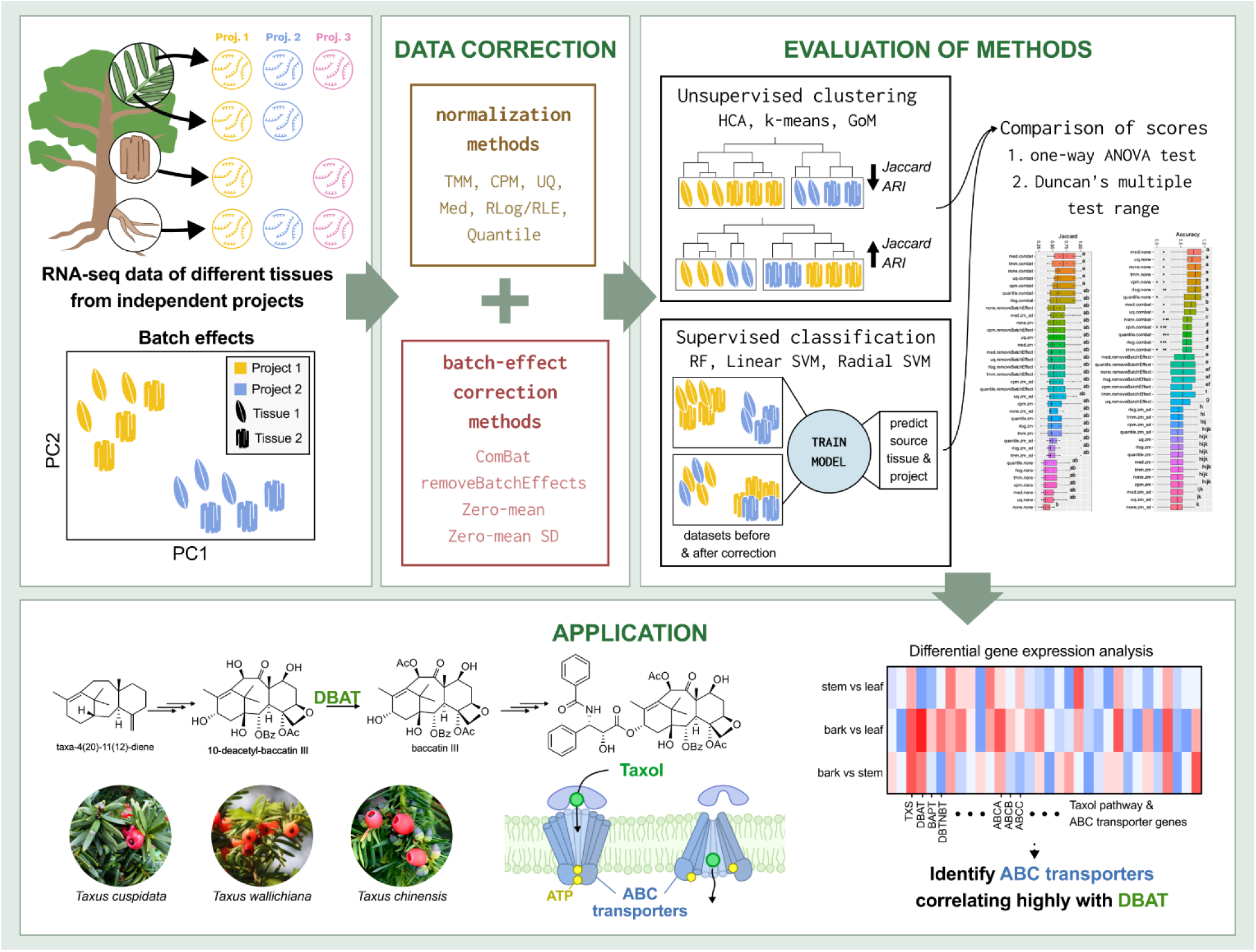

## 1. Introduction

The yew family (Taxaceae) is a diverse group of evergreen trees and shrubs found throughout the world. This coniferous family is made up of two extinct and six extant genera, *Amentotaxus*, *Austrotaxus*, *Cephalotaxus*, *Pseudotaxus*, *Torreya*, and *Taxus*, each with its unique characteristics and distribution patterns^1^. The genus *Taxus* benefits from a widespread distribution across the northern hemisphere with 12 natural species, *T. floridana* Nutt., *T. brevifolia* Nutt., *T. chinensis* Rehd., *T. contorta* Griff., *T. wallichiana* Zucc., *T. globosa* Schlecht., *T. mairei* Lem., *T. calcicola* L.M. Gao, *T. canadensis* Marshall, *T. cuspidata* Zucc., *T. baccata* L., and *T. florinii* Spjut. These are followed by two hybrids, *T. media* Rehder and *T. hunnewelliana* Rehder^2^. They are valued not only for their ornamental appearance^3^, but also for their long history of medicinal use, due to the production of chemotherapeutic agents paclitaxel (Taxol) and related taxanes^4–7^. To date, over 550 taxanes have been discovered and characterized, of which taxol and its related precursors, baccatin III (BAC III) and 10-deacetylbaccatin III (10-DAB III), are of the utmost importance in cancer treatment and research^8^. The anticancer compounds were initially extracted from the inner bark of *T. brevifolia* (1963). However, the indiscriminate harvesting of the trees compounded with the slow-growing nature of *Taxus* plants has brought the species to “Near Threatened” status and led to critical environmental concerns^2^. Fortunately, over the last two decades, alternatives for obtaining taxol have emerged, including plant cell cultures, semi-synthesis from its precursors^9^, chemical total synthesis^10^, use of endophytic fungi as a source^11^, and biosynthetic production using metabolically engineered heterologous hosts^12,13^, such as *E. coli*^14^, yeast^15,16^, and tobacco^17,18^.

Like many living organisms, gene expression in the genus *Taxus* follows a dynamic pattern, which is governed by factors such as species, environmental cues, plant tissue types, and sampling date. In this sense, RNA sequencing (RNA-seq) has emerged as a highly accurate and cost-effective approach to analyze expression differences (transcriptomic studies) between various species of the *Taxus* genus. The first such transcriptomic study was reported from cultured cells of *T. cuspidata* in 2010^19^, and additional ones were gradually added. This technique has made it possible to address challenging issues related to the detection of differentially expressed genes (DEGs), discovery of novel enzymes, characterization of metabolic pathways, and so on. Over the past decade, several transcriptomic studies have contributed to our understanding of taxol biosynthesis, however, they also revealed inconsistencies and gaps. For example, Hao et al. reported tissue-specific transcriptomes of *T. mairei* and analyzed the expression of candidate taxol biosynthetic genes across root, stem, and leaf tissues^20^. Similarly, Yu et al. compared *T. media* and *T. mairei* and suggested that variation in taxoid content could be attributed to differential expression of pathway genes^21^. However, different studies often report distinct sets of candidate genes for the same enzymatic steps, raising uncertainty about which genes are truly functional in vivo.

Beyond genes involved in biosynthetic pathways, new transcriptional regulators of the pathway have also been proposed. Li et al. demonstrated that elicitation with methyl jasmonate (MeJA) activates TF families such as MYB, bHLH, ERF, AP2, and MYC^22^, while Zhang et al. identified a large set of WRKY transcription factors from *T. chinensis*, among which TcWRKY8 and TcWRKY47 were shown to enhance the expression of multiple taxol-biosynthesis-related genes^23^. Despite these findings, not all studies agree on the identity or functional relevance of specific TFs. In parallel, miRNA-mediated^24^ control of terpenoid backbone and paclitaxel biosynthesis has been uncovered in cultured *Taxus × media* cells, further emphasizing the complexity of multilayered regulation. At the genomic level^25^, evidence of whole-genome duplication (WGD) has further complicated pathway reconstruction. Xiong et al. identified multiple highly similar gene copies within taxol-related clusters, such as two *TS* genes (TS2 and TS3, 99.96% identical) and two *T5αH* genes (98.67% identical)^26^, raising questions about functional redundancy versus specialization.

Despite its remarkable advantages, RNA-seq data (whether as bulk or single cell) generally suffer from a phenomenon called “*batch effect*”. This term refers to systematic dissimilarities between samples that are not attributable to experimental design, and that originate instead from personnel differences, experimental procedures, operators, sampling time disparities, as well as variations in batches of reagents, and even biological deviations, such as growth location, to name a few common sources^27–29^. These effects appear in many bio-based research areas, including high-throughput metabolomics^29^ and high-throughput transcriptomics, and interfere with downstream statistical analyses^27–29^. Despite efforts to mitigate their occurrence through proper experimental protocols and design, they may persist and prove difficult to rectify. Thus, it is imperative to exercise caution and meticulousness when working with batched data in order to ensure the success and reproducibility of research.

In RNA-seq-based data assays, various strategies are employed depending on the objectives of a particular project. For example, researchers may investigate gene expression patterns between different species, tissues under similar or differing environmental conditions, or ontogenic stages. In this context, the *Taxus* genus has seen several publicly available RNA-seq datasets published by different research groups, encompassing multiple species, conditions, and tissues. However, collecting data from multiple sources can present challenges due to the potential for batch effects, which introduce heterogeneity into the data. In recent years, several studies have attempted to address the issue, exploring the effects of normalization methods and between-sample normalization using batch-effect correction (BEC) on gene expression analysis^30–33^. The accurate identification of transcriptomic differences through read counts depends on initial steps of read mapping to the reference genome and their subsequent quantification^34^ of transcriptomics features, such as genes, exons, or isoforms ^34–36^. Prior to downstream differential expression (DE) analysis, it is critical to perform preprocessing steps such as the filtering of lowly expressed genes^37^, normalization^31,38,39^, and BEC^27,34,40,41^. Filtering removes low-abundance genes to reduce false discovery rates^34^. Although normalization was once consider optional for RNA-seq ^42^, it has since been established as critical step for accurate interpretation^43^. Indeed, the choice of normalization method often influences DE results more than the statistical testing approach^30^. This necessity arises from the variable proportion of mRNA attributed to particular genes across biological conditions^43^, driven by factors such as environmental stimuli, disease states, and developmental stages. Without proper normalization, comparisons risk reflecting differences in overall mRNA composition rather than true biological variation. Therefore, normalization remains an essential step in RNA-seq pre-processing pipelines, correcting for technical factors such as sequencing depth, RNA composition, and gene length^34^.

Taken together, these studies highlight both progress and persisting ambiguities in understanding taxol biosynthesis. Different research groups have reported varying candidate genes for the same pathway steps, multiple layers of regulation by distinct TF families, and gene duplications that confound the differentiation between functional paralogs and artifacts of redundancy. To resolve these issues, the integration of transcriptomic datasets is necessary. However, such integrative analyses are strongly confounded by batch effects across independent studies. Batch effect correction (BEC) therefore becomes an essential prerequisite to harmonize datasets, minimize technical noise, and enable reliable discovery of consistent pathway components. Despite substantial advances, the genetic and regulatory basis of taxol biosynthesis, and more broadly taxane metabolism, remains incompletely understood, with critical uncertainties yet to be resolved.

BEC studies have primarily focused on human or animal RNA-seq data, leaving a gap in understanding the impact of these methods on plant gene expression patterns. In order to bridge this gap, our current study applied a total of 35 different combinations of normalization and BEC techniques to a meta-dataset developed from various independent investigations, species, and tissues, in order to identify the most effective approach for batch effect removal. This comprehensive and rigorous approach (Fig. 1A) allowed us to investigate the effects of these methods not only on whole *Taxus* transcriptome, but also on key genes involved in taxol biosynthesis and potential ATP-binding cassette (ABC) transporters. Previous studies have demonstrated that samples from different batches display greater variation than those from different tissues processed within the same batch, indicating that gene expression patterns may distinguish batches more readily than tissue types (Fig. 1B). However, following BEC, it is expected that samples from the same tissue will be more accurately grouped together (Fig. 1C). Therefore, the implementation of BEC techniques is crucial for ensuring the accuracy and reliability of downstream analyses, such as the identification of differentially expressed genes, in the genus *Taxus*. This important research sheds light on the best practices for analyzing plant RNA-seq data and will ultimately contribute to a more accurate understanding of complex gene expression in plants, particularly in the context of taxol biosynthesis and transport. With this airtight analysis, we aim to pave the way for more in-depth studies on the impact of normalization and BEC methods in the field of plant biology.

**Figure 1.**
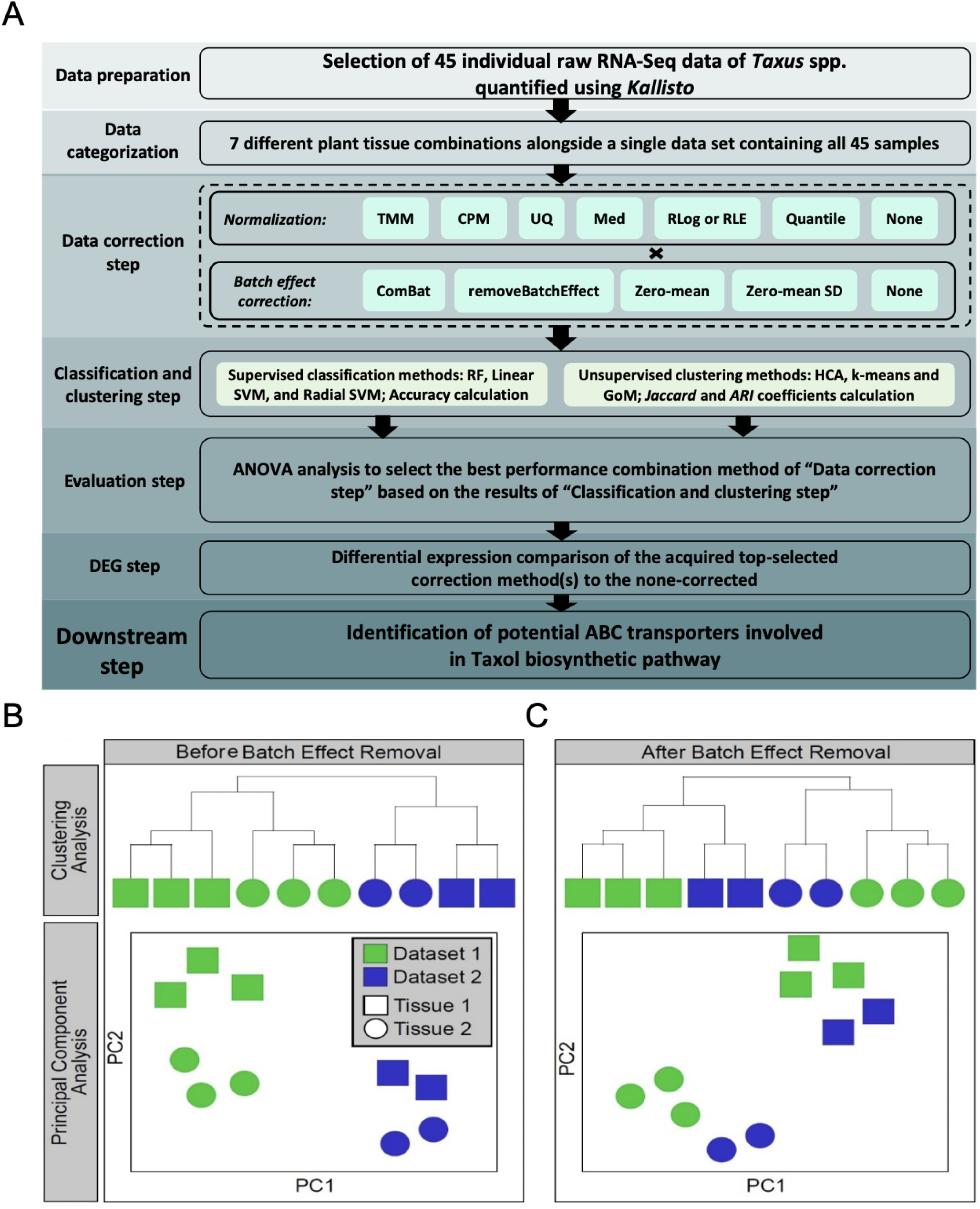
Schematic representation of the bioinformatic pipeline and rationale for batch effect correction. **A.** The steps undertaken in this study to investigate efficient methods for preprocessing RNA-seq data are presented. **B** and **C** show the mechanism of batch effect removal in two independent projects with two different tissues is illustrated using two prevalent unsupervised clustering methods: hierarchical clustering (HCA) (upper panel) and principal component analysis (PCA) (lower panel). As shown, prior to BEC, the samples were clearly separated based on the corresponding datasets 1 and 2 (**B**), whereas application of BEC grouped the samples from the independent datasets by tissue type (**C**).

## 2. Materials and Methods

### 2.1. Gene expression dataset collection

All RNA-seq data of *Taxus* spp. were retrieved from SRA database. To do this, the database was searched for RNA-seq datasets harboring more than three samples, and subsequently, only the datasets with more than one tissue in common were selected. This resulted in three independent datasets (PRJNA661543, PRJNA493167, and PRJNA730337) from three *Taxus* species (*T. wallichiana*, *T. cuspidata*, and *T. chinensis*) and four different tissues (Bark, Leaf, Root, and Stem) resulting in 45 samples (Table 1). For each RNA-seq sample, the FASTQ files were downloaded using fastq-dump (from SRA Toolkit version 3.0.7), the reference genome and the annotation for *T. chinensis* (GCA_019776745.2_Ta-2021_genomic.fna and GCA_019776745.2_Ta-2021_genomic.gtf, respectively) were retrieved from NCBI. The FASTQ files were trimmed using Trimmomatic^44^ version 0.39 (with parameters `SE - phred33 input.fastq.gz output.fastq.gz ILLUMINACLIP:TruSeq3-SE.fa:2:30:10 LEADING:3 TRAILING:3 SLIDINGWINDOW:4:15 MINLEN:36`), and subsequently kallisto^45^ (version 0.44.0) was applied to trimmed FASTQ files for quantification of gene expression.

**Table 1.** Datasets used in the study.

| Project ID | *Taxus* species | Plant tissues | Number of RNA-seq samples |
| --- | --- | --- | --- |
| PRJNA661543 | *T. wallichiana* | Bark | 6 |
|  |  | Leaf | 3 |
|  |  | Root | 3 |
|  |  | Stem | 3 |
| PRJNA493167 | *T. cuspidata* | Leaf | 4 |
|  |  | Root | 4 |
|  |  | Stem | 4 |
| PRJNA730337 | *T. chinensis* | Bark | 6 |
|  |  | Leaf | 6 |
|  |  | Root | 6 |
| **Total** | 3 | - | 45 |

### 2.2. Data Partitioning for normalization and batch effect

As mentioned above, prior to applying BEC, when analyzing the data using principal component analysis (PCA) and/or hierarchical clustering analysis (HCA), it is expected that samples from the same batch (i.e., project) will cluster together. Meanwhile, for batch effect-free data, we expect that samples from the same tissue (e.g., leaf) will be gathered in the same cluster (Fig. 1). To test this hypothesis, we constructed different combined datasets. We first combined samples from projects that shared the same tissues, for all available combinations of tissues. For example, the tissues bark and leaf were present in both projects PRJNA661543 and PRJNA730337, while leaf tissue was not present in project PRJNA493167. Therefore, we created a combined dataset of all the bark and leaf samples from projects PRJNA661543 and PRJNA730337, but not PRJNA493167. We created an additional dataset that combines all 45 samples. The details of all these combinations are shown in Table 2.

**Table 2.** Details of eight tissue combinations developed from three independent projects.

|  |  | PRJNA661543 | PRJNA493167 | PRJNA730337 | Total |
| --- | --- | --- | --- | --- | --- |
| **Bark-Leaf** | Bark | 6 | - | 6 | 21 |
|  | Leaf | 3 | - | 6 |  |
| **Bark-Leaf-Root** | Bark | 6 | - | 6 | 30 |
|  | Leaf | 3 | - | 6 |  |
|  | Root | 3 | - | 6 |  |
| **Bark-Root** | Bark | 6 | - | 6 | 21 |
|  | Root | 3 | - | 6 |  |
| **Leaf-Root** | Leaf | 3 | 4 | 6 | 26 |
|  | Root | 3 | 4 | 6 |  |
| **Leaf-Root-Stem** | Leaf | 3 | 4 | - | 21 |
|  | Root | 3 | 4 | - |  |
|  | Stem | 3 | 4 | - |  |
| **Leaf-Stem** | Leaf | 3 | 4 | - | 14 |
|  | Stem | 3 | 4 | - |  |
| **Root-Stem** | Root | 3 | 4 | - | 14 |
|  | Stem | 3 | 4 | - |  |
| **All** | Bark | 6 | - | 6 | 45 |
|  | Leaf | 3 | 4 | 6 |  |
|  | Root | 3 | 4 | 6 |  |
|  | Stem | 3 | 4 | - |  |

### 2.3. Normalization and batch effect-correction approaches

To apply normalization and remove batch effects, a pipeline similar to the one described by Vandenbon (2022) was applied. Six normalization and four batch-effect-removal approaches were combined. Including a run without normalization and one without batch effect removal, we have 35 combinations in total. For each combined dataset, normalization was applied, the data was log-transformed, and batch effect removal was implemented. The normalization and batch effect approaches are summarized as below.

#### 2.3.1. Normalization methods

In RNA-seq experiments, factors such as sequencing depth will cause between-sample variations, leading to challenges in downstream analyses. To adjust these between-sample biases, a number of different normalization approaches have been proposed. The following six normalization approaches were applied in this study to assess the effect of normalization on the analysis.

*Trimmed Mean of M-values* (*TMM*) estimates the ratio of RNA productions between two samples, based on the assumption that the number of differentially expressed genes is small. To this end, the method compares a test sample to a reference sample, excludes genes with the highest expression levels and large log ratios, and then calculates the weighted average of the remaining log ratios. If the RNA production is similar between the two samples, this average should be close to 1; otherwise, it provides a scaling factor for normalizing the test sample^46^. TMM was performed using the `calcNormFactors` function in *edgeR* package^47^ (version 3.42.4).

##### Counts per million (CPM)

In the field of data analysis, counts per million (CPM) is a widely used method for normalizing data to make accurate comparisons among different samples. The method takes into consideration the total number of counts in each sample by dividing the count value of each data point by the total count in that sample. The resulting ratio represents the relative abundance of a particular data point in a given sample. Multiplying this ratio by 1 million yields the CPM value i which ensures the sum of all CPM values in a sample will equal 1 million. The approach allows for a standardized comparison of data across varying sample sizes, ensuring that one sample with a higher total count does not skew the results ^48^.

##### Upper Quartile (UQ)

The UQ calculates the upper quartile of a sample (75^th^ percentile) after excluding the zero values, which may skew results and inaccurately reflect the true nature of the data. This exclusion ensures that the resulting UQ value is a more accurate representation of the sample’s performance. The expression value of each gene in each sample is divided by the UQ of that sample, then each sample is multiplied by the average UQ of all samples^30,31^.

*Median* (*Med*) is similar to UQ, except that it calculates the median of non-zero values when calculating the normalization factor^31^.

*Regularized Logarithm* (*RLog*) is similar to log-transformation, however, for genes with low counts, the values of different samples will shrink together^49^. The *DESeq2* package (version 1.40.2) was used to perform this normalization^49^.

*Quantile normalization* works by making the quantiles of all samples similar such that the value at a given percentile is the same for all samples. The quantile normalization was performed using `normalizeQuantiles` function from the *limma* package^50^ (version 3.56.2).

#### 2.3.2. Batch effect removal approaches

Combat from the *sva* package^51^ (version 3.48.0) followed by `removeBatchEffect` from the *limma* package^50,52^ were developed to remove non-biological variations introduced into data. In both cases, by specifying the batch covariate (in this study, the source dataset) and biological covariate (in this study, the tissues), these functions will remove batch effects while keeping biologically meaningful variations in the data.

*Zero-mean per batch* (or mean-centering; abbreviated as *zm*) and *Zero-mean and standardized per batch* (abbreviated as *zm_sd*) are simple approaches that have been previously employed for removing batch effects from expression data^53^. In the first approach, the mean expression value of each gene within a batch is subtracted from all samples in that batch, resulting in a zero mean for each gene. In the second approach, in addition to subtracting the mean, the expression values of each gene within a batch are divided by their standard deviation, resulting in a unit standard deviation.

### 2.4. Unsupervised clustering methods and supervised classifiers

#### 2.4.1. Unsupervised clustering methods

We applied three different unsupervised clustering methods, *k*-means clustering, hierarchical clustering analysis (HCA), and Grade of Membership models (GoM) (https://proceedings.mlr.press/v22/taddy12.html)^54^ for clustering of biological samples of *Taxus* RNA sequencing data. The GoM models are statistical methods that utilize the concept of partial membership to classify observations based on their likelihood of belonging to a particular group. This method takes into account the different degrees of association that individuals may have with a particular group, rather than simply assigning a binary label of membership or non-membership. The resulting classification is a more nuanced representation of individuals within a population, as it allows for the possibility of individuals belonging to multiple groups to varying degrees. This approach has been applied in a variety of fields such as genetics, psychology, and market research, and has proven to be a powerful tool for understanding complex data. Overall, the use of GoM models offers a more comprehensive approach for classification, enabling a deeper understanding of complex data and its underlying patterns.

For *k*-means clustering we used the `kmeans` function (with default parameters and `nstart`=10) from the *stats* package (v4.3.1) in R. For HCA, we used Euclidean distance and average linkage (using `dist`, `hclust`, and `cutree` functions from the *stats* package (v4.3.1) in R). For GoM, we used the `topics` function from the *maptpx* package (v1.9-7) in R. We ran the topics function on the `exp` of the normalized and batch corrected data (because the normalized data is already log transformed, and `topics` function expects positive numbers). The gene expression heatmaps showing the column clustering results were made using the *ComplexHeatmap* package (v2.16.0) in R.

We applied these clustering methods to samples in each of the eight combined datasets (Table 2), after applying the 35 normalization and batch effect-removal methods. When necessary, the clustering was performed twice: once with the number of clusters set to the number of tissues, and once with the number of clusters set to the number of source datasets. We then asked how similar the resulting clusters are to tissue groups and dataset groups (clustering results where the number of clusters were set to the number of tissues will be compared to tissue groups, and clustering results where the number of clusters were set to the number of datasets will be compared to dataset groups). To measure similarity of cluster assignments and the sample groups, we used the *Jaccard index*^55^ and the *Adjusted Rand Index* (*ARI*)^56^. Briefly, for each cluster, we found the group with the highest *Jaccard index* to that cluster, and then calculated the weighted average of these values over all clusters (weighted by cluster size):

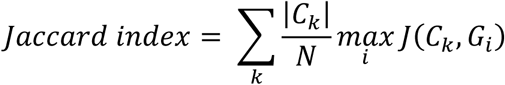

where *C_k_* is *k*^th^ cluster, *G_i_* is the *i*^th^ group (*i*^th^ tissue or *i*^th^ dataset), *N* is the number of samples in the combined dataset, and *J* is the *Jaccard index*, defined as 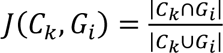. For each normalization and batch effect removal scenarios, this procedure will result in 8 values when comparing the tissue groups and 8 values when comparing the dataset groups (one value per combined dataset).

In addition to the *Jaccard index*, we also calculated the *ARI* to measure similarity between the cluster assignments and the true labels (tissue groups and dataset groups). The *ARI* is calculated as:

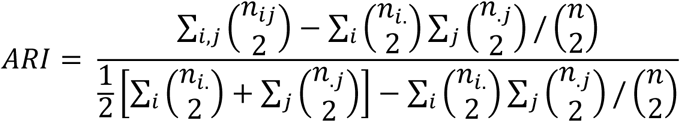

where *n_ij_* is the number of samples belonging to cluster *i* and group *j* (for example *tissue_j_* or *dataset_j_*) and *ni*. is the number samples in cluster *i* and *ni* is the number of samples in group *j*. To calculate *ARI*, we used `adjustedRandIndex` from the *mclust* package (v6.0.0) in R. To assess how different normalization and batch effect-removal scenarios perform, we applied the one-way analysis of variance (ANOVA) test to these values (using `aov` from the *stats* package (v4.3.1) in R), followed by the Duncan multiple range test (*p* < 0.05) for mean comparison (using `duncan.test` from the *agricolae* package (v1.3.7) in R).

#### 2.4.2. Supervised machine learning models

Another way to assess the performance of normalization and batch effect-removal methods, is to test the following two key questions: 1) “Can tissue identity be better predicted after batch effect removal?” and 2) “Can the source dataset of each sample be better predicted before batch effect removal?”. To assess these two questions, we developed two experiments. First, we used the expression value of 100 most highly expressed genes as features for the classification task, and next, we applied PCA (`prcomp` from the *stats* package (v4.3.1) in R) on the data and used the first 10 PCs as features for the classification. For classification task, we used Support Vector Machine (`svm` from the *e1071* package, with radial or linear kernel), and Random Forest (*randomForest* package in R). For evaluation of the classification, we employed “*hold-out validation*” (also known as “split sample method”), in which the dataset is split into two non-overlapping parts of “*training*” and “*test*” datasets. Here, 50% hold-out validation was used, in which the data is partitioned into two equal parts, and the model is trained on the first part and tested on the other ^57^. This process was repeated 100 times with random partitions, resulting in 100 test accuracy values for each task (prediction of tissue or source dataset), and for each classification method. This approach was applied to each of the 8 combined datasets and 35 normalization and batch effect removal scenarios. To assess performance under different scenarios, for a given task (prediction of tissue or source dataset) and a given classification method (Random Forest, or SVM), we combined the accuracy values from the 8 combined datasets and performed a one-way ANOVA test, followed by Duncan’s multiple range test (*p* < 0.05) for mean comparison.

### 2.5. Differential expression analysis

To evaluate the effect of normalization and batch effect-removal methods, we examined the similarity of the resulting differentially expressed genes (DEGs) across the combined datasets. Briefly, for each combined datasets, after applying normalization, log-transformation, and batch effect removal, we used the *limma* package^50^ to identify DEGs for each pair of tissues. Genes with the adjusted *p*-value ≤ 0.05 were considered DEGs, and the DEG sets were compared between all the pairs of the 35 normalization and batch removal scenarios by calculating the number of genes in common (overlap) between the gene sets of the two methods, and furthermore, by calculating the *Jaccard index* between the gene sets of the two methods. We then calculated the average number of overlaps, and average of the *Jaccard index*, for each pair of scenarios over all pairs of tissues.

### 2.6. Identification of Gene Ontology terms for *Taxus* genes

The DNA sequence of genes (i.e., sequences of predicted transcripts) were extracted from reference genome in the FASTA format using GffRead v0.12 ^58^. The extracted DNA sequences were subsequently aligned to the RefSeq non-redundant proteins (NR) using `blastx` (version 2.15.0). The blast2go tool ^59^ was used to assign Gene Ontology (GO) terms to the *Taxus* genes using these `blastx` results (Supplementary Data 1, Supplementary Table 1).

### 2.7. Selecting paclitaxel biosynthesis related genes

The coordinates of the genes related to paclitaxel biosynthetic pathway were retrieved from the supplementary Table 23 of ^60^. The coordinates of two additional genes (TOT1 and T9aH1) were retrieved from ^61^. The *T. chinensis* (GCA_019776745.2 Ta-2021) GTF file was searched, and any genes overlapping these coordinates were selected. Genes located more than 10,000 bp distance from the reported coordinates were excluded. This resulted in 25 genes, reported in Supplementary Data 1, Supplementary Table 2.

### 2.8. Selecting ABC transporter genes

To identify transporter genes, we selected all genes associated with related GO terms (“ATP-binding cassette (ABC) transporter complex”, “ABC-type manganese transporter activity”, “ABC-type protein transporter activity”, and “ABC-type transporter activity”), which resulted in 225 genes. Moreover, we retrieved protein sequence of 167 ABC transporters in *Arabidopsis thaliana* from JGI’s Phytozome portal (https://phytozome.jgi.doe.gov/). These proteins were aligned to the DNA sequences of *Taxus* genes using `tblastn`, and genes with more than 70% identical positions were selected, resulting in 47 genes (34 of which overlapped with the GO list). The DNA sequence of these genes were translated into amino acid sequences using the `getorf` program from *EMBOSS* package (version 6.6.0.0). We then scanned these sequences with hmmscan from the HMMER package (version 3.3.2) and Pfam-A HMMs from InterPro database. Sequences containing the ABC transporter-related domains were further submitted to SMART website, and sequences with confirmed ABC-related domains were selected. This resulted in selecting 189 genes out of the original 238 genes candidates (reported in Supplementary Data 1, Supplementary Table 3).

### 2.9 Structural modelling and molecular docking

To evaluate whether the selected ABC transporter candidates could accommodate taxane intermediates, molecular docking was performed using 10-deacetylbaccatin III (10-DAB III) as a representative substrate. Candidate transporters selected from the transcriptomic correlation analysis were structurally modelled using AlphaFold-predicted protein structures^62^. The human taxol-associated ABC transporter structure PDB ID: 9CTF was used as a control template to define the potential substrate-binding region^63^.

The transporter models were aligned to the 9CTF template, and the ligand-binding cavity coordinates from the template were transferred to the corresponding region of each *Taxus* candidate. Molecular docking was carried out using AutoDock Vina^64^. For each transporter candidate, 10-DAB III was docked into the predicted central cavity with box size 30 Angstrom to allow comparison across candidates. Docking poses were ranked according to predicted binding affinity, reported in kcal/mol.

## 3. Results

### 3.1. Unsupervised clustering methods

To investigate the impact of normalization and BEC on sample grouping, we assessed clustering and PCA patterns before and after applying the rlog+ComBat method to the combined tissue dataset. Supplementary Fig. 1A-B illustrates PCA plots of combined dataset Bark+Leaf, before and after normalization and batch effect removal using rlog+ComBat. Similarly, *k*-means clustering (k = 2) and Grade of Membership (GoM) clustering (k = 2) on the same dataset are presented in Supplementary Fig. 2A-B and Supplementary Fig. 3A-B, respectively. Prior to normalization and BEC, samples grouped mostly by project, whereas post-correction clustering aligned with tissue type. To systematically evaluate 35 normalization and batch effect removal combinations, we performed hierarchical clustering, *k*-means, and GoM analyses across all 8 tissue combinations described in Table 2. Cluster assignments were compared against tissue and dataset labels for each sample using the *ARI* and the *Jaccard index* (Fig. 2-3, Supplementary Fig. 4, see Methods).

**Figure. 2.**
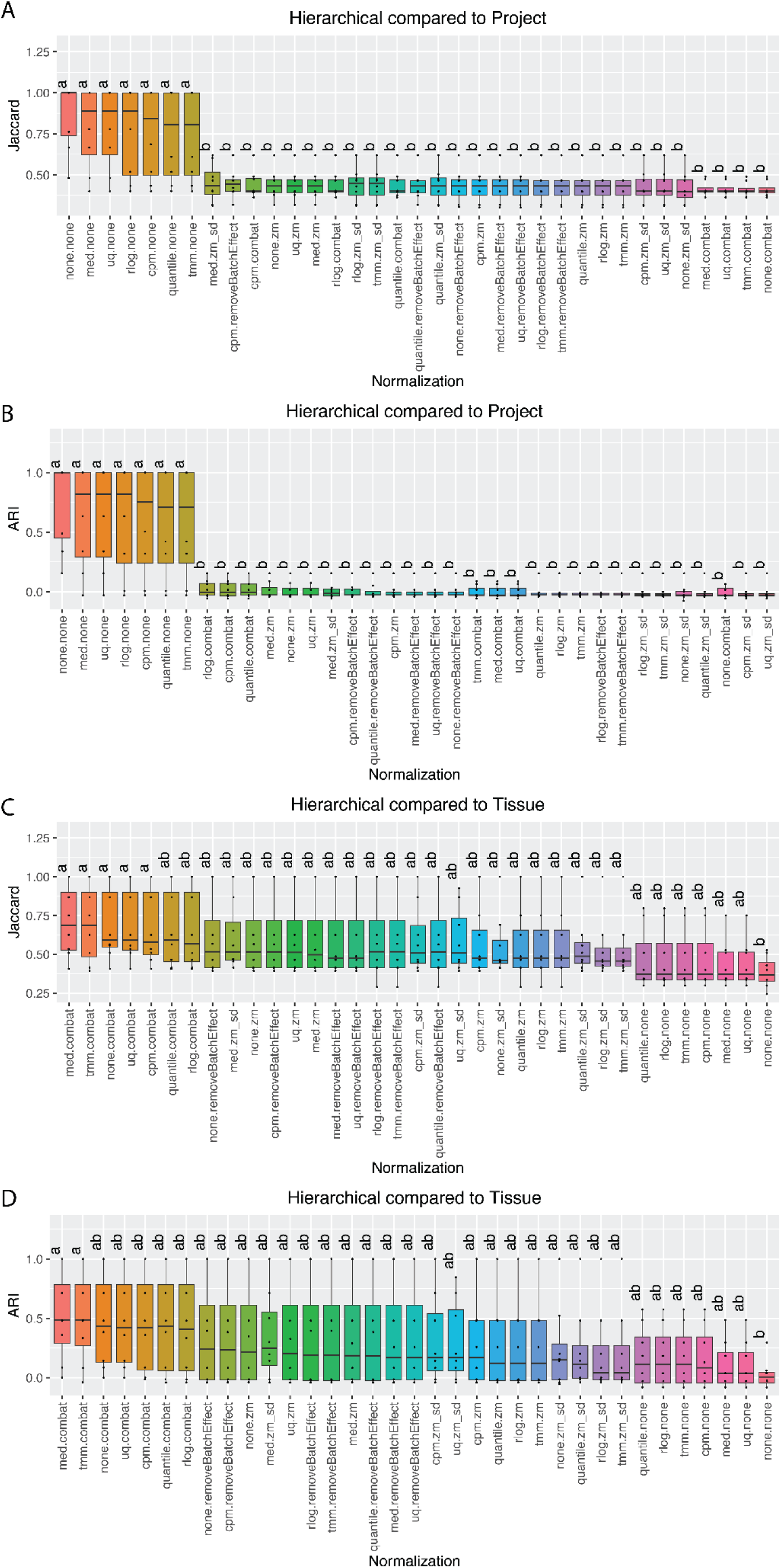
Comparison of 35 different combinations using the box plot-assisted Duncan multiple range test for the HCA clustering method. Combinations consisted of 7 normalization methods combined with 5 BEC approaches. Test results, with significance determined at *p* < 0.05 threshold, were based on either the *Jaccard index* or *ARI*, with each box in the box plots containing 8 values. **A** and **B** represent comparisons of HCA cluster assignments to the project IDs (dataset of origin) using the *Jaccard index* and *ARI*, respectively. **C** and **D** represent comparisons with “tissue types” using the same indices.

**Figure 3.**
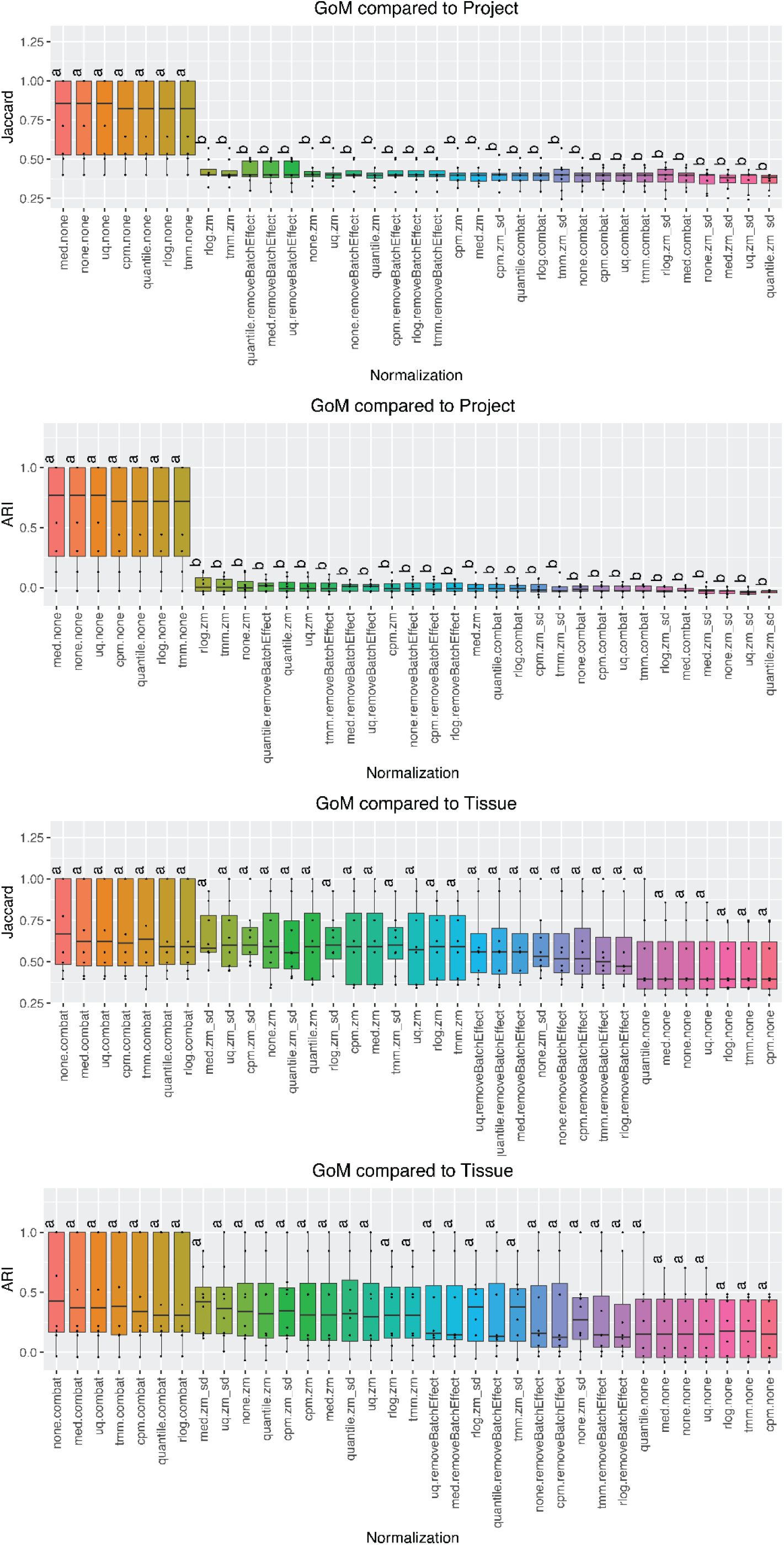
Comparison of 35 different combinations using the box plot-assisted Duncan multiple range test for the GoM clustering method. The combinations consist of 7 normalization methods multiplied by 5 BEC approaches. Test results with significance determined at *p* < 0.05, are based on either the *Jaccard index* or *ARI*, with each box in the box plots containing 8 values. **A** and **B** represent the comparison of GoM cluster assignments to the project IDs (dataset of origin) using the *Jaccard index* and *ARI*, respectively. **C** and **D** show comparisons of GoM cluster assignments to tissue types using the same metrics.

We expected that RNA-seq samples prior to BEC to cluster primarily by batch (dataset of origin). ANOVA results confirmed significant differences among all the 35 correction methods in terms of *Jaccard index* across the eight tissues combinations. When comparing the HCA cluster assignments to project IDs using Duncan mean comparison, the 35 methods separated into two groups (Fig. 2A-B). Group a, consisting of the seven no-batch-correction methods (none.none, med.none, uq.none, rlog.none, cpm.none, quantile.none, and tmm.none) exhibited the highest *Jaccard index* values. The remaining 28 combinations were classified in group b with lower *Jaccard index* scores. A similar grouping was observed when clustering was compared to tissues labels (Fig. 2C-D). ComBat-corrected methods (med.combat, tmm.combat, none.combat, uq.combat, and cpm.combat), achieved the highest *Jaccard index* values (group a), while none.none (no normalization and no batch effect removal) produced the lowest *Jaccard index* values (group b). *ARI* values followed the same trend, with med.combat and tmm.combat in group a, and none.none in group b (Fig. 2D).

To confirm the robustness of our findings, we also applied *k*-means clustering (Supplementary Fig. S4). The results were consistent with HCA analysis, showing that methods without batch correction clustered by project but performed poorly by tissue, while combined normalization and batch correction achieved the highest *Jaccard index* and *ARI* scores. This reinforces the necessity of batch-effect correction with normalization for accurate tissue classification.

We further applied GoM model to allow partial sample membership and provide a more flexible clustering framework. When comparing clusters to project IDs, GoM analysis revealed significant differences between methods with and without batch effect removal. All no-batch-correction combinations clustered in category a (Fig. 3A-B). When examining clusters by tissue type, ComBat batch-corrected combinations exhibited higher *Jaccard index* and *ARI* scores, whereas those without batch correction showed the lowest values. However, no significant differences were observed among the 35 combinations, with all placed in category a (Fig. 3C-D). These findings demonstrate the presence of batch effect across RNA-seq datasets and underscore the significance of batch effect removal. Together, normalization and BEC enhance detection of underlying biological variation, ensuring accurate tissue classification independent of variability among *Taxus* species, project origin, or other factors.

### 3.2. Supervised classification approaches

To assess the performance of different normalization and BEC methods, we evaluated how accurately a sample’s tissue type or source dataset could be predicted from its gene expression values. Higher accuracy in predicting the source dataset (project IDs) indicates the presence of batch effect, whereas higher accuracy in predicting the tissue type or biological trait suggests successful batch-effect removal. We used either the expression value of 100 most highly expressed genes, or the first 10 principal components of the entire expression matrix to train various classifiers for predicting sample tissue type or source dataset (see Methods). ANOVA results revealed significant differences among the 35 correction methods in terms of accuracy values across all eight tissues combinations.

Using Random Forest classifiers trained on principal components (PCs) of gene expression data to predict project IDs (source dataset of each sample), significant differences were observed among the normalization and batch effect-removal methods. Duncan’s mean comparison test classified the 35 correction methods into 15 groups (a-o groups) (Fig. 4A). The highest prediction accuracies (group a) were observed for five methods: cpm+none, none+none, rlog+none, tmm+none, and quantile+none (Fig. 4A). Consistent patterns were observed using alternative classification algorithms or when using the expression values of the 100 most highly expressed genes rather than the top 10 PCs (Table 3; Fig. 4B-F). Notably, methods incorporating zero-mean normalization and standardization for BEC consistently underperform. Overall, BEC methods exhibited reduced accuracy relative to uncorrected methods in predicting project IDs, reflecting effective removal of batch-associated signal.

**Figure 4.**
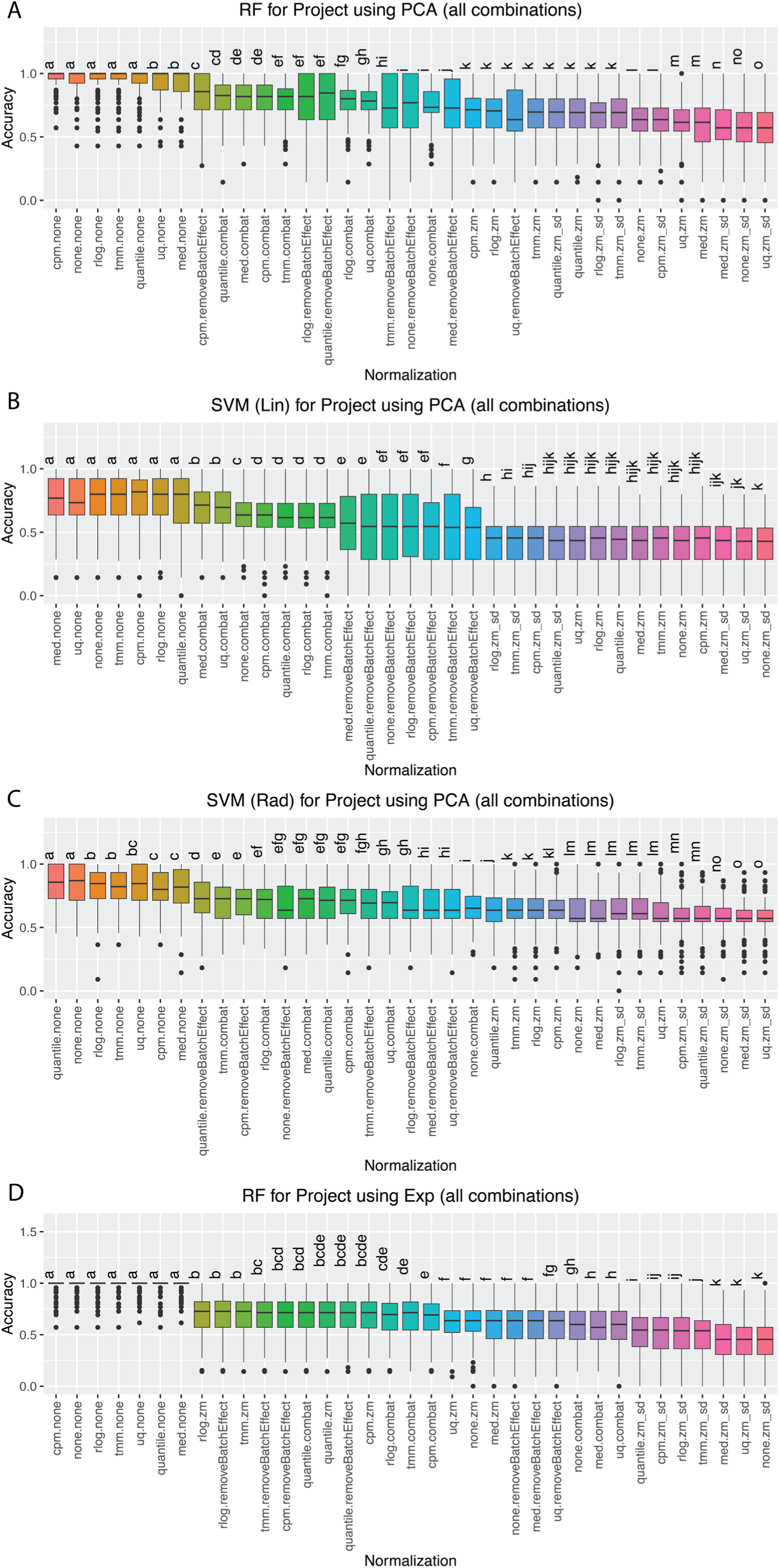
Performance comparison of 35 normalization and BEC methods for predicting sample source datasets, based on Duncan’s multiple range test (*p* < 0.05). Each box included 800 accuracy values (100 train/test splits for 8 tissue combinations; see Methods). RF, SVM (Lin), and SVM (Rad) classification of project IDs using the top 10 principal components (all combinations) are shown in panels **A-C**, respectively, while classification using the top 100 most highly expressed genes (all combinations) is shown in panels **D-F**, respectively.

**Table 3.** Methods with the highest accuracy when predicting Project IDs and tissue type.

| **Classifiers** | **Training Data** | **Top methods when predicting Project IDs** | **Top methods when predicting Tissues** |
| --- | --- | --- | --- |
| **Random Forest** | 10 PCs | cpm+none, none+none, rlog+none, tmm+none, and quantile+none | cpm+combat, rlog.combat, uq.combat, tmm.combat, none.combat, and quantile.combat |
|  | 100 Genes | cpm.none, rlog.none, tmm.none, uq.none, quantile.none, med.none, and none.none | rlog.combat, quantile.combat, tmm.combat, and cpm.combat |
| **SVM (Linear)** | 10 PCs | med.none, uq.none, tmm.none, cpm.none, rlog.none, quantile.none, and none.none | med+removeBatchEffect and uq+removeBatchEffect |
|  | 100 Genes | med.none, uq.none, tmm.none, cpm.none, rlog.none, quantile.none, and none.none | rlog.combat, tmm.combat, cpm.combat, quantile.combat, and med.combat |
| **SVM (Radial)** | 10 PCs | quantile.none, none.none, rlog.none, tmm.none, uq.none, cpm.none, and med.none | uq.none, med.none, none.none, and tmm.none |
|  | 100 Genes | med.none, uq.none, tmm.none, cpm.none, rlog.none, quantile.none, and none.none | rlog.combat, tmm.combat, cpm.combat, and quantile.combat |

When using Random Forest to predict tissue labels, the 35 combined methods were divided into 12 clusters (a-l). Methods using ComBat as batch effect removal (cpm+combat, rlog.combat, uq.combat, tmm.combat, none.combat, and quantile.combat) were among the best performers (Fig. 5A). Similar trends were observed when using other classification methods and when using the expression values of top 100 most expressed genes instead of top 10 PCs, where methods with batch effect removal performed best, while method without batch effect removal were among the lowest performing methods (Table 3, Fig. 5B-F). An exception to this trend was observed with SVM using a radial kernel applied to 10 PCs of expression data, where surprisingly, methods with no batch effect removal (uq.none, med.none, none.none, and tmm.none) were among the best-performing approaches (Table 3).

**Figure 5.**
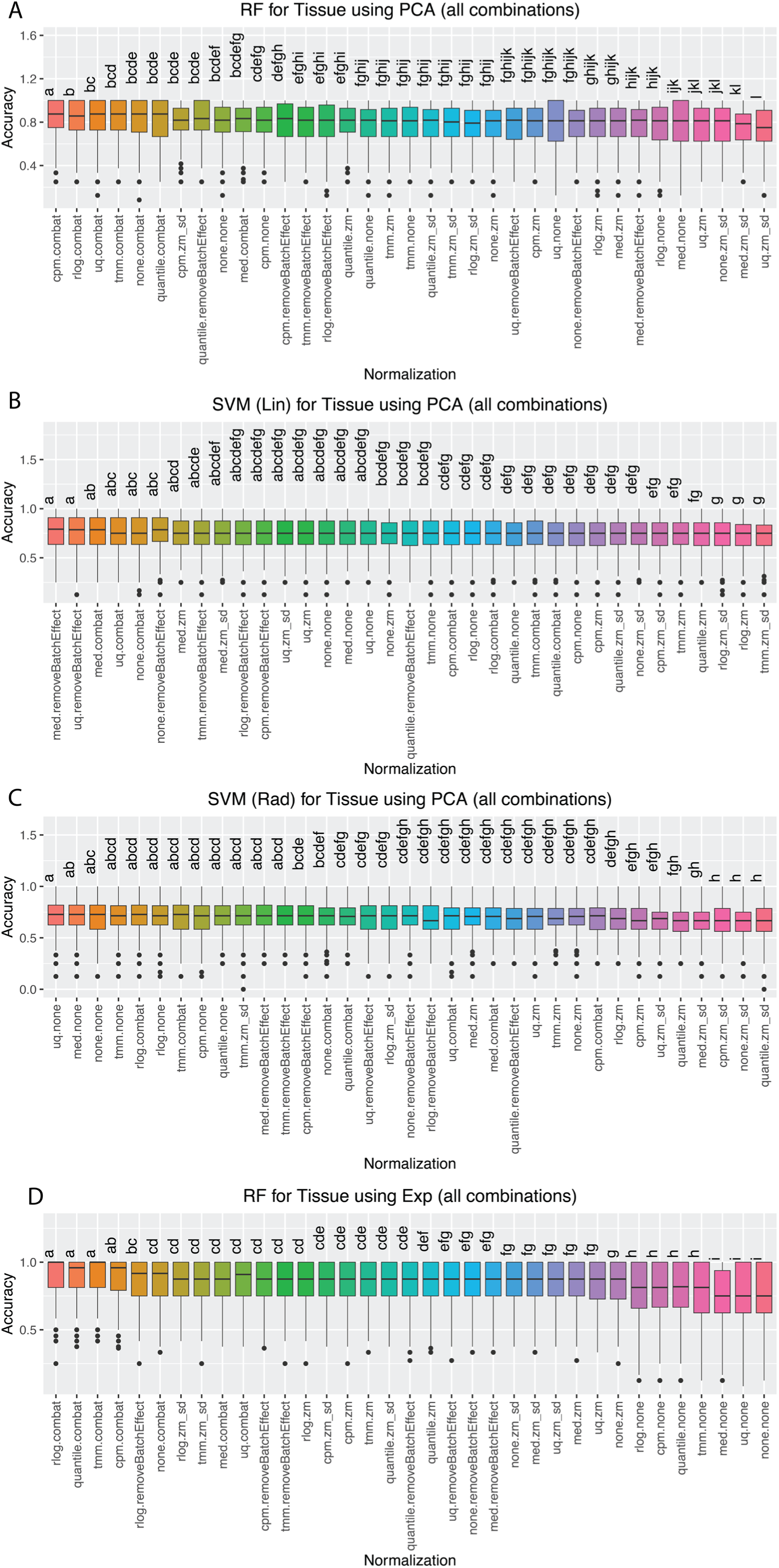
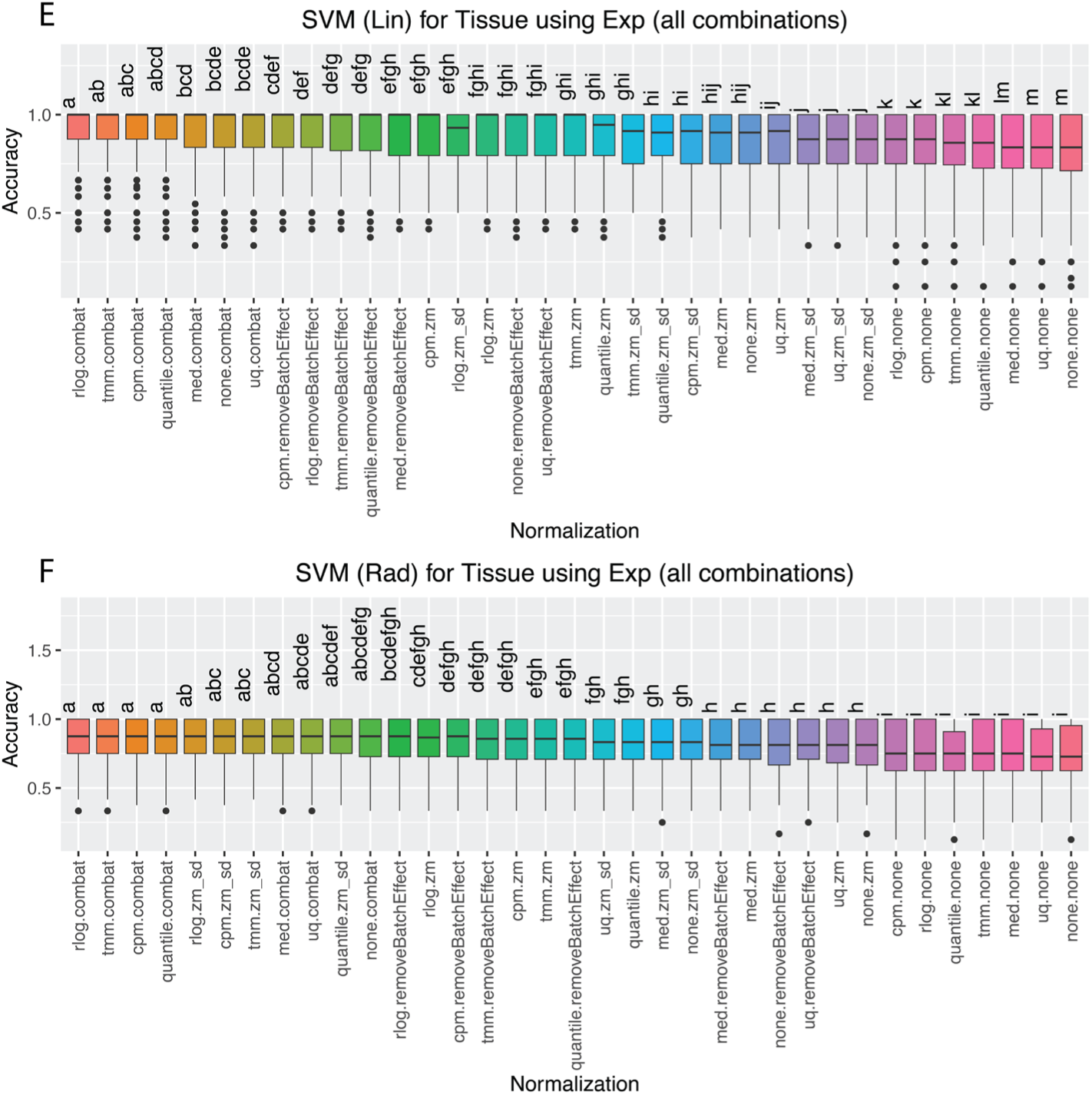
Performance comparison of 35 different normalization and BEC methods for predicting sample tissue, based on Duncan’s multiple range test (*p* < 0.05). Each box includes 800 prediction accuracy values (100 train/test splits for 8 tissue combinations, see Methods). RF, SVM (Lin), and SVM (Rad) sample tissue classification using the top 10 principal components (all combinations) are shown in panels **A–C**, respectively, and classification using the top 100 most highly expressed genes (all combinations) is shown in panels **D–F**, respectively.

### 3.3. Differential expression analysis

To assess the effect of normalization and batch effect removal on downstream investigations such as functional studies, we performed differential expression analysis on the normalized, log-transformed, and batch corrected gene expression data. Fig. 6A shows the average overlap in the number of differentially expressed genes (DEGs) between pairs of normalization and batch effect methods, averaged across tissues pairs from the combined datasets. As expected, we observed significant differences before and after BEC. Quantile normalization, rlog, cpm, and tmm generated more similar results to each other irrespective of the batch effect method used, whereas the zero-meaning and standardizing batch effect removal method appear to yield less similar results. Among all the 35 methods, quantile.combat, rlog.combat, cpm.combat, and tmm.combat exhibited the highest number of overlaping DEGs. Notably, these four combinations also ranked among the best performing methods in terms of tissue prediction accuracy from gene expression values and concordance between clustering results and true labels. Fig. 6B shows the average *Jaccard index* of DEG overlap between pairs of normalization and batch effect methods. Consistent with the DEG counts, methods without batch effect removal have low similarity to those with batch effect removal. Furthermore, regardless of the BEC method, quantile, rlog, cpm, and tmm normalization produced results that were more similar to each other than to those from other normalization methods.

**Figure 6.**
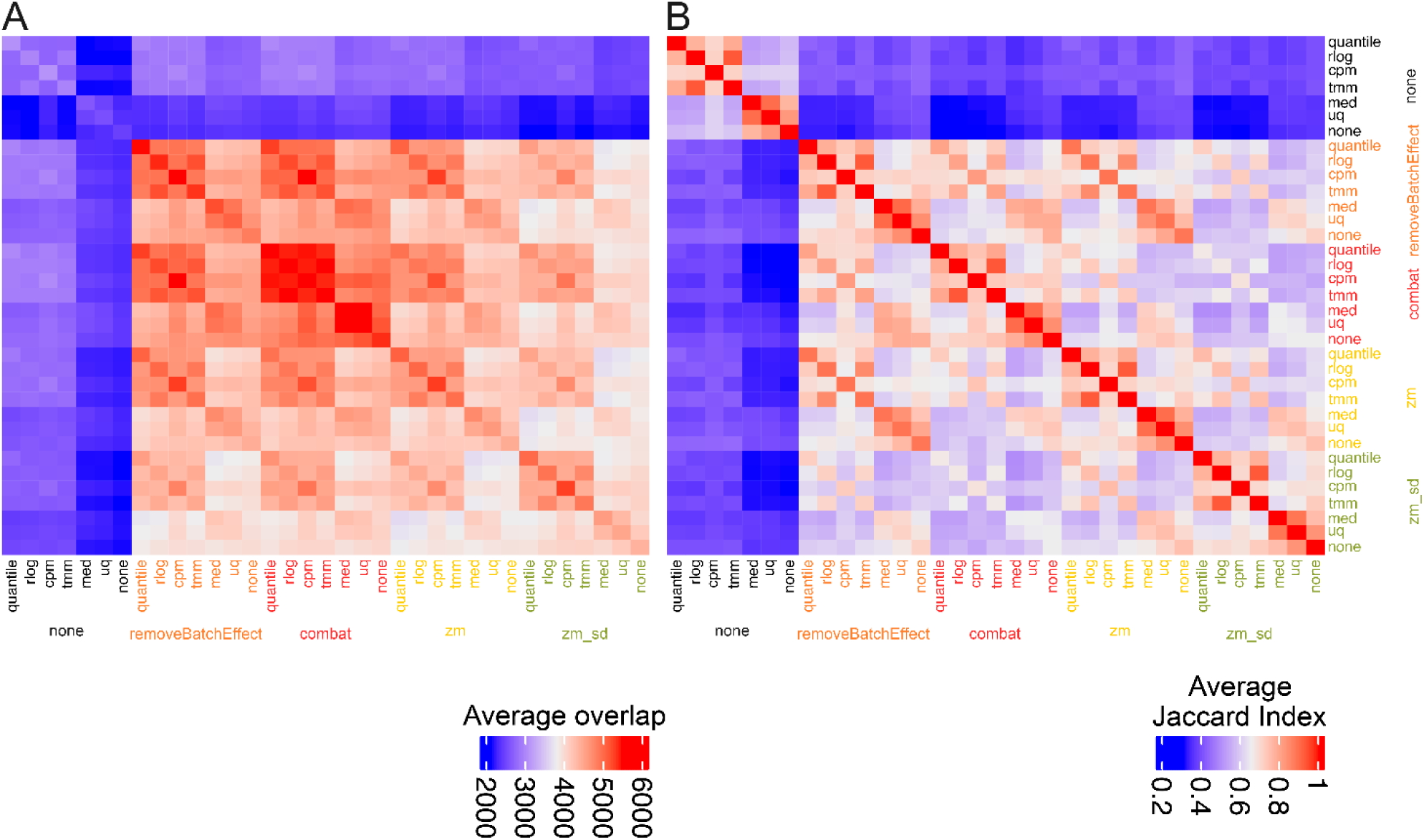
Average number of differentially expressed genes (DEGs) between pairs of normalization and batch effect methods (averaged over different tissue pairs from the combined datasets. **A**. Average number of DEGs, **B**. Average *Jaccard index*.

### 3.4. Extraction of the ABC transporters in the genome of Taxus genus

To identify possible ABC transporters available in the *Taxus* genome, a systematic bioinformatic analysis was applied (see Methods). A total of 189 putative ABC transporters were identified across four tissues: stem, root, bark, and leaf (Supplementary Table 3). Analyses focused on the optimal preprocessing pipeline, combining regularized log transformation and ComBat batch effect correction (rlog.combat), applied to the complete four-tissue dataset (bark, leaf, stem, root; Supplementary Fig. 5). These ABC transporter candidates were clustered into two main categories, clearly distinguished by high and low differential expression levels (Supplementary Fig. 5). Subsequent correlation analyses were performed to explore associations between these transporters and genes involved in Taxol biosynthesis, using a heatmap clustered dendrogram generated from the rlog.combat-processed dataset for the combination of all four tissues (Fig. 7A). Notably, 11 of the 189 transporters were excluded due to invariant expression profiles, precluding meaningful analyses, as correlation scores are undefined for constant vectors.

**Figure 7.**
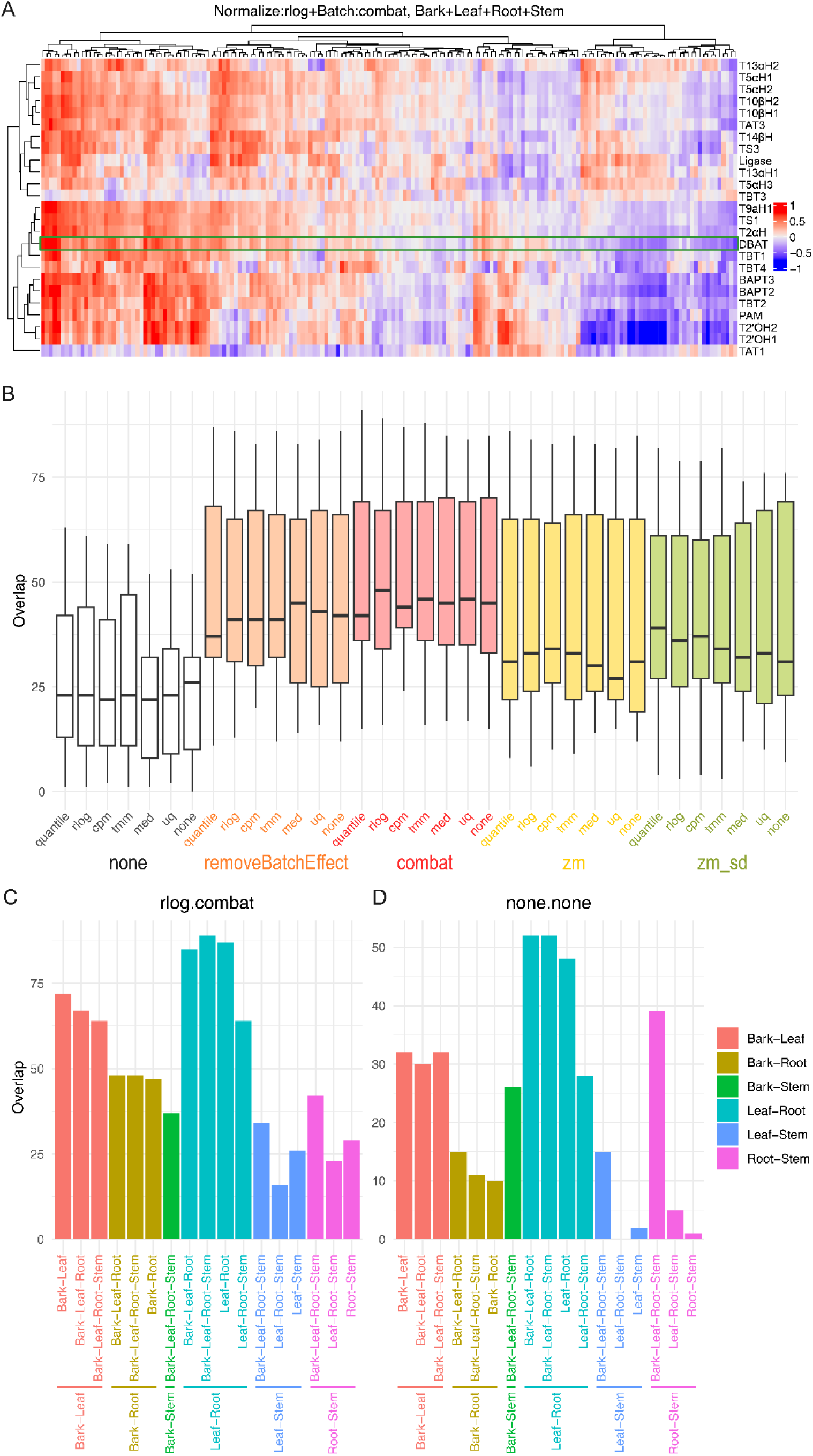
Correlation Analysis and Gene Distribution of ABC Transporters and Paclitaxel Biosynthesis Genes Across Tissue Combinations and Preprocessing Methods. **A.** Clustered heatmap depicting correlations between putative ABC transporters and genes involved in the taxol biosynthetic pathway. The dendrogram was constructed using the optimal preprocessing pipeline (rlog normalization combined with ComBat batch effect correction) applied to data from all four tissues (bark, leaf, stem, and root). Number of ABC transporters and paclitaxel biosynthesis-related genes among the DEGs are shown for **B**. each combination of normalization and BEC method (n = 17; each box corresponds to number of genes for different tissue combinations), and in different tissue combinations for **C**. rlog.combat and **D**. none.none.

Given the central role of the 10-deacetylbaccatin-III-10β-O-acetyltransferase (DBAT) gene in the Taxol biosynthetic pathway (Supplementary Fig. 6), we focused on ABC transporters exhibiting strong correlation with the expression level of this gene. We identified six top candidates (Supplementary Table 4), which were further characterized via BLASTx and Conserved Domain Database searches (https://www.ncbi.nlm.nih.gov/Structure/cdd/wrpsb.cgi) to determine their family classification. The first (rna-gnl|WGS:JAHRHJ|evm2.model.Chr04.1261) and fourth (rna-gnl|WGS:JAHRHJ|evm2.model.Chr08.2758) highly expressed ABC transporter genes, were assigned to the ABC transporter B family and named *Tc*ABCB1 and *Tc*ABCB4. The third candidate (rna-gnl|WGS:JAHRHJ|evm2.model.Chr01.3318) was classified within the ABC transporter C family and named *Tc*ABCC3, whereas the second (rna-gnl|WGS:JAHRHJ|evm2.model.Chr01.3803), fifth (rna-gnl|WGS:JAHRHJ|evm2.model.Chr05.1210) and sixth (rna-gnl|WGS:JAHRHJ|evm2.model.Chr05.1206) highly expressed ABC transporter genes belonged to the ABC transporter G family, and named *Tc*ABCG2, *Tc*ABCG5, *Tc*ABCG6.

### 3.5. Extraction of the number of taxol biosynthetic genes among DEGs of each combined method

To further assess the effect of normalization and batch effect removal methods on differential expression analysis, we examined the number of ABC transporter and paclitaxel biosynthesis-related genes among DEGs for each tissue pair combination (Table 2) after applying these methods (Fig. 7B). As expected, the number of these genes increased after BEC, though counts of such genes varied across tissue pair and combinations. Fig. 7B-C show the counts of ABC transporters and paclitaxel biosynthesis-related genes among DEGs for rlog.combat and none.none, respectively. Rlog.combat exhibited greater overlap with DEGs compared to none.none. Notably, we observed an increased overlap for Leaf *vs*. Root and Bark *vs*. Leaf pairs compared to Bark *vs*. Stem, Leaf *vs*. Stem, or Root *vs*. Stem DEGs (Fig. 7C), a pattern that was also seen, to a lesser extent, with none.none (Fig. 7D). Detailed number for DEGs and associated gene counts per method and tissue pair, are reported in Supplementary Data 2, Supplementary Table 5. These findings suggest that these transporters may play important roles in *Taxus* species, potentially involved in the localization, transport, or secretion of taxol or its intermediates within the plant.

### 3.6. Molecular docking supports the potential interaction of predicted ABC transporters with 10-DAB III

To further evaluate whether the DBAT-correlated ABC transporter candidates could plausibly interact with late-stage taxane intermediates, we performed structure-based docking using 10-deacetylbaccatin III (10-DAB III) as a representative substrate. Because the selected candidates were predicted to belong to ABC transporter families commonly associated with small-molecule export, the transporters were modelled as multidrug exporters. AlphaFold-predicted structures of the top transporter candidates as well as a human ABCB1 (PDB ID: 9CTF) as a control model were inspected in the context of an inward-facing exporter conformation^63^. In this state, the transmembrane domains form a central cavity that is open toward the cytosolic side, consistent with the expected substrate-loading state of ABC exporters.

Docking of 10-DAB III revealed that several top candidates could position the ligand within the central cavity of the transmembrane region with sufficient binding affinity. The docked substrates were generally located in the inward-facing cavity, surrounded by transmembrane helices (Fig. 8). Among the tested candidates, *Tc*ABCC3 showed the most favorable predicted affinity, with a docking score of −8.38 kcal/mol, followed by *Tc*ABCG2 with −7.57 kcal/mol (Table 4). These values were more favorable than those observed for the control models, which showed affinities of −6.37 and −6.89 kcal/mol, respectively. *Tc*ABCB4 and *Tc*ABCB1 showed intermediate affinities of −6.78 and −6.49 kcal/mol, while *Tc*ABCG5 displayed a weaker predicted affinity of −6.21 kcal/mol. These results suggest that the candidate transporters not only accommodate 10-DAB III in the central cavity but may also provide binding environments more compatible with the ligand than the control models.

**Figure 8.**
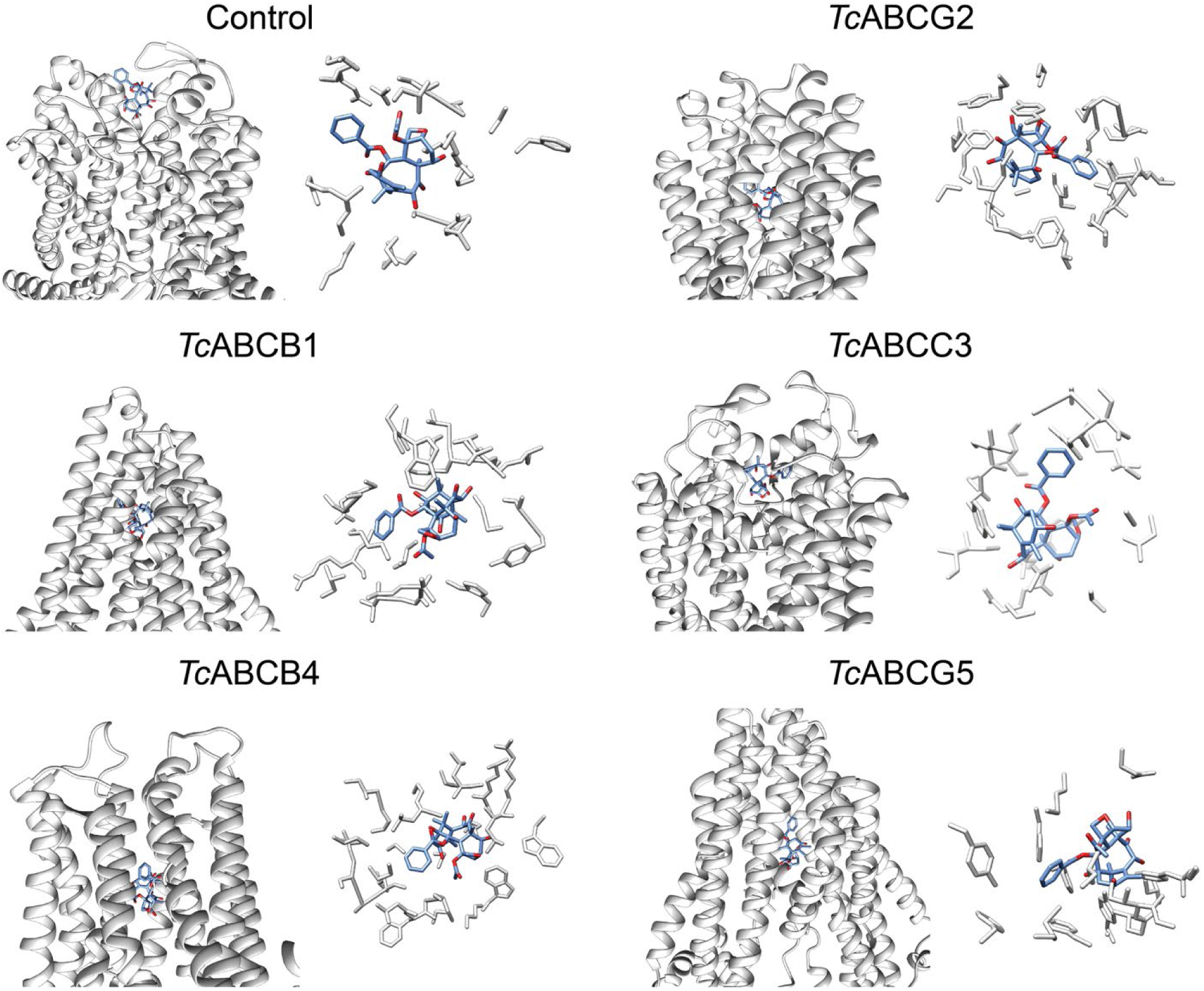
Structural modelling and molecular docking of 10-deacetylbaccatin III with top predicted ABC transporter candidates and human ABCB1 (PDB ID: 9CTF) as control. For each model, the left panel shows the docked ligand positioned within the predicted central transmembrane cavity, while the right panel highlights residues located within 5 Å of the ligand.

**Table 4.** Predicted binding affinities of 10-deacetylbaccatin III docked to top ABC transporter candidates.

| Model ID | Affinity (kcal/mol) | Intramolecular Energy (kcal/mol) |
| --- | --- | --- |
| Control (Human ABCB1) | -6.16 | 8.92 |
| TcABCB1 | -7.08 | 1.05 |
| TcABCG2 | -6.44 | 1.15 |
| TcABCC3 | -8.38 | 7.11 |
| TcABCB4 | -6.78 | 0.67 |
| TcABCG5 | -6.21 | 0.80 |

Visual inspection of the docking poses further supported this interpretation. In the prioritized candidates, 10-DAB III was positioned within or near the central transmembrane cavity, surrounded by residues lining the predicted substrate-binding region. This cavity location is consistent with an inward-facing multidrug exporter state, where hydrophobic and partially polar substrates are first captured from the cytosolic side of the membrane before conformational transition toward an outward-facing state^65^. The observed docking poses therefore provide structure-based support for the hypothesis that selected *Taxus* ABC transporters may participate in the export or intracellular redistribution of taxane intermediates. Notably, the docking-based ranking showed consistency with the transcriptome-derived ranking of candidate transporters.

Candidates that were highly ranked based on their expression correlation with DBAT also tended to show favorable docking behavior with 10-DAB III, including improved predicted affinity relative to the control models. This agreement between independent transcriptomic and structure-based analyses strengthens the selection of these transporters, suggesting that the candidates most closely associated with taxol biosynthetic gene expression are also structurally compatible with binding a late-stage taxane intermediate. In particular, the overlap between the DBAT-correlation ranking and docking performance supports the use of an integrated candidate-selection strategy, combining co-expression and substrate-binding affinity to identify ABC transporters for functional validation. Further experimental validation is required to confirm their functions.

## 4. Discussion

Despite significant advancements in single-cell RNA sequencing (scRNA-seq) and spatial transcriptomics, bulk RNA-seq remains a fundamental and widely used method in biological research^66^. Its versatility and experimental flexibility make it a valuable complement to scRNA-seq approaches. In certain cases, the homogeneity of cell populations justifies averaging, while generating single-cell or single-nucleus suspensions can be challenging or unfeasible. Furthermore, biases towards specific cell types in scRNA-seq data may limit its overall interpretability. The high per-sample costs associated with scRNA-seq constrain the number of replicates and conditions that can be studied^66^. The implementation of bulk RNA-seq technology has greatly contributed to acquiring critical biological insights, and is broadly recognized worldwide^34,66^. This acceptance can be attributed to its ability to measure the expression levels across large gene sets simultaneously, providing a comprehensive view of complex biological systems. By enabling high-throughput sequencing of RNA transcripts from heterogenous cell populations, bulk RNA-seq has transformed our understanding of gene expression patterns, cellular pathways, and molecular mechanisms underlying various biological processes. In plants, bulk RNA-seq, together with whole-genome sequencing, have been particularly instrumental in elucidating biosynthetic pathways involved in the biotechnological production of plant secondary metabolites^67–70^. By analyzing the abundance and expression levels of RNA transcripts, this powerful technology has revealed potential mechanisms and key enzymes responsible for the synthesis of these important compounds. Its far-reaching applications continue to expand and contribute to the further advancement of biological research across multiple fields.

Batch effects are a pervasive issue that can significantly impact the integrity and reproducibility of high-throughput data^40,41^. It is therefore crucial to understand the factors influencing the effectiveness of BEC. One key consideration is the degree of confounding between class and batch factors. When these factors overlap substantially, correcting batch effects without introducing bias in the class factor becomes challenging. Furthermore, the normalization method applied during data preprocessing can affect BEC success, as different approaches may yield varying results and can potentially introduce or mask batch effects. Lastly, the characteristics of the batch effect itself, including its magnitude and variability, also influence the performance of correction methods. Recognizing these factors is essential for selecting the most appropriate BEC approach and ensuring accurate and reproducible analyses in high-throughput studies.

### Upstream normalization procedures alone or combined with unsupervised clustering-based BEC methods

To evaluate the impact of upstream normalization procedures on BEC analysis in *Taxus* genus, six normalization approaches were applied. Two scenarios were considered. The first focused on three popular unsupervised clustering methods, hierarchical clustering, *k*-means, and GoM, and assessed their performance using the *Jaccard index* and *ARI*. When comparing normalization-only approaches to the none.none combination, no statistically significant differences were observed. The none-none combination, followed by normalization-only approaches, yielded the highest values for both metrics, whereas applying BEC methods alone or in combination with normalization produced the lowest values of both metrics. Higher metric values when clustering the source dataset (i.e., project IDs) suggest the presence of batch effect in the current three datasets.

Analysis based on tissue type revealed that the combined use of normalization and BEC methods is dependent on the clustering method and similarity indexes utilized. For hierarchical clustering and *k*- means, the highest *Jaccard index* and *ARI* values were achieved with combined normalization and BEC methods. Specifically, the ComBat method produced the highest *Jaccard index* values in hierarchical clustering, while med.combat and tmm.combat combinations yielded the highest *ARI* values. For *k*-means clustering, the highest *Jaccard index* values were observed for rlog.combat, rlog.removeBatchEffect, tmm.removeBatchEffect, uq.zm, tmm.combat, and quantile.combat combinations, whereas both tmm.removeBatchEffect and rlog.removeBatchEffect combinations yielded the maximum *ARI* values. The relatively high values when clustering bytissue (i.e., biological trait) suggest successful batch effect removal across datasets. For the GoM method, however, none of the 35 combinations produced significant differencesin either metric, as all samples formed a single cluster. It is worth noting that hierarchical clustering, *k*-means and the GoM model serve as effective dimension reduction techniques and hold potential for mitigating batch effects ^27,71^. Nevertheless, the absence of apparent batch effects in Fig. 2C and 2D is not fully understood and is consistent with the previous observations^54^. The data’s specific characteristics and the chosen dimension reduction methods likely influence the results. A deeper analysis and evaluation of the variables at play is necessary to fully understand of effective batch effect control and to guide the development of more refined and optimized dimension reduction techniques for mitigating batch effects in future studies.

### Upstream normalization procedures alone or combined with supervised classification-based BEC methods

In addition to evaluating the effectiveness of normalization and BEC methods through traditional metrics, their impact was further examined on two specific factors: the ability to accurately predict the tissue of each sample after batch effect removal, and the ability to correctly predict the source dataset of each sample before batch effect removal. To assess these factors, we first utilized the expression values of the 100 most highly expressed genes as features for classification. Additionally, we applied PCA and utilized the first 10 PCs as features for classification. Evaluating classification performance for accurate tissue type and source dataset prediction provided a comprehensive understanding of their effectiveness in minimizing batch effects and normalizing data.

When applying three supervised classification algorithms to predict project IDs using either the top 10 PCs or the top 100 most expressed genes, the highest accuracy was achieved with “none-none” combination as well as normalization-only approaches, despite some observed fluctuations. Duncan’s test results identified the quantile.none most frequently as yielding maximum accuracy. In contrast, the use of BEC methods, either alone or combined with normalization, resulted in lower accuracy for project ID prediction, suggesting the presence of batch effects among the three datasets.

For tissue label prediction, normalization combined with BEC generally produced the highest accuracy values, indicating effective batch effects removal across datasets. However, the best-performing combination among the 35 tested varied by classification algorithm, suggesting that accuracy depends strongly on the type of supervised classification algorithm. Notably, an exception was SVM with radial kernel applied to 10 PCs, where, surprisingly, methods with no batch effect removal (uq.none, med.none, none.none, and tmm.none) performed among the best. Overall med.removeBatchEffect, cpm.combat (RF) and uq.removeBatchEffect (SVM Lin) were the most effective combination for batch effect removal using the top 10 PCs. In contrast, using the top 100 most expressed genes, rlog.combat consistently achieved high accuracy across all three supervised classifiers, suggesting strong BEC in these datasets.

Considering all data derived from clustering and classification analyses, upstream normalization alone did not provide sufficient performance in removing batch effect in the current dataset, which is consistent with previous findings^38^. Similar outcomes were observed when applying BEC methods alone. However, combinations of normalization with BEC methods exhibited different rankings, with only some yielded better results. The observation that upstream normalization does not have a substantial influence on performance suggests that this factor may play a limited role in explaining discrepancies among BEC rankings reported in the literature^38^. Variations in performance among BEC methods may simply be attributed to the unique characteristics of the datasets being studied. Alternatively, the rankings may reflect minor distinctions, emphasizing the potential for inconsistency and lack of generalizability in these results^38^. Therefore, although upstream normalization may not be a significant contributing factor, further research is required to identify determinants of BEC effectiveness. Furthermore, the context and limitations of each individual study should be carefully considered.

### Structural analysis of taxane-binding ABC transporters

Although much of taxol pathway discovery has focused on biosynthetic enzymes, the transport and subcellular distribution of taxane intermediates are likely to represent an additional layer of pathway regulation. Taxanes are complex hydrophobic diterpenoids, and their accumulation within membranes or the cytosol may influence cellular homeostasis, enzyme accessibility, and pathway flux^72^. Heterologous production of taxol intermediates in yeast, tobacco, or other chassis is often limited not only by enzyme activity and precursor supply, but also by product toxicity, intracellular retention, and inefficient recovery of hydrophobic intermediates. Taxus-derived exporters could therefore provide useful engineering modules to enhance taxane secretion, reduce intracellular stress, and improve product extraction.

Incorporating transporter discovery into taxol biosynthesis studies may help shift pathway engineering from enzyme reconstruction alone toward a more complete system that includes biosynthesis, compartmentalization, and metabolite trafficking. Therefore, the identification of ABC transporters co-expressed with late taxol biosynthetic genes provides important insight into how *Taxus* cells may coordinate biosynthesis with metabolite trafficking. In this study, several ABC transporter candidates showing strong expression correlation with DBAT also exhibited sufficient binding affinity toward 10-deacetylbaccatin III, suggesting that these transporters may be transcriptionally coordinated with late-stage taxane metabolism. The agreement between the docking-based ranking with the transcriptome-derived ranking strengthens the identification of these candidates for downstream functional validation.

## 5. Conclusions

In this study, we systematically evaluated preprocessing correction methods and found that upstream normalization alone is insufficient for effective batch effect removal in *Taxus* RNA-seq datasets. Although normalization independently yielded high accuracy in predicting dataset origin, this performance primarily reflects the persistence of batch effects rather than authentic biological signals. By contrast, combining normalization and BEC enabled the identification of genuine biological variation within the data, exemplified by the distinct tissue types analyzed, thereby demonstrating more effective batch effect mitigation. Nevertheless, a universal combination strategy is unrealistic as the optimal method depends on the choice of classification algorithm and feature selection, highlighting the inherent complexity and dataset-specific nature of BEC.

Our results demonstrated the presence of batch effect across the RNA-seq samples obtained from disparate datasets and underscore the need of integrating normalization with appropriate BEC to improve data comparability and biological insight in transcriptomic studies. Future research should focus on refining these strategies and identify factors driving variability in BEC performance towards establishing more generalizable best practices for RNA-seq data analysis in plants. Using this framework, we identified several ABC transporter candidates showing strong correlation with DBAT, a key enzyme involved in the conversion of 10-deacetylbaccatin III to baccatin III. Molecular docking further supported the biological plausibility of these candidates, as several prioritized transporters were predicted to accommodate 10-deacetylbaccatin III within the central transmembrane cavity of inward-facing multidrug exporter models. The agreement between transcriptomic ranking and docking-based prioritization strengthens the hypothesis that these transporters may participate in taxane intermediate recognition, trafficking, or export. Our findings are poised to enhance the accuracy of downstream analyses, thereby facilitating robust investigations into the molecular mechanisms underlying specialized metabolism in *Taxus* species.

## Supplementary Materials

Supplementary Information: Supplementary Figures 1-6. Supplementary Data 1: Supplementary Tables 1-4. Supplementary Data 2: Supplementary Table 5.

## Acknowledgements

This work was financially supported by the Canadian Institutes of Health Research (CIHR) Grant (202203PJT-481041-TIR-CFAA-98996), and New Frontiers in Research Fund (NFRF) – Exploration Grant (2022-00158) to C.I. and Y.X., Y.D received funding from McGill Engineering International Tuition Award (MEITA) and Fonds de recherche du Québec - Santé (FRQS) Master’s Research Scholarships (351750). C.X received funding from FRQNT Master’s Research Scholarships (351047) and McGill Engineering Undergraduate Student Master’s Award (MEUSMA).

## CRediT author statement

**Jaber Nasiri:** Conceptual Design, Data Curation, Investigation, Methodology, Formal Analysis, Validation, Writing – original draft. **Alireza Fotuhi Siahpirani:** Bioinformatics analysis, Data Curation, Investigation, Methodology, Formal Analysis, Validation. **Yueming Dong:** Structural analysis, Investigation, Methodology, Formal Analysis, Validation, Visualization and Writing - review & editing. **Catherine Xu:** Methodology, Visualization and Writing - review & editing. **Yu Xia:** Writing - review & editing. **Codruta Ignea:** Conceptualization, Funding Acquisition, Supervision, Visualization, Writing – review & editing.

## References

1 Lange, B. M. & Conner, C. F. Taxanes and taxoids of the genus Taxus–A comprehensive inventory of chemical diversity. Phytochemistry 190, 112829 (2021).

2 Thakur, A. & Kanwal, K. S. Assessing the Global Distribution and Conservation Status of the Taxus Genus: An Overview. *Trees*, Forests and People, 100501 (2024).

3 DeLong, J. M. & Prange, R. K. Taxus spp.: Botany, Horticulture, and Source of Anti-Cancer Compounds. Horticultural Reviews 32, 299–327 (2006).

4 Weaver, B. A. How Taxol/paclitaxel kills cancer cells. Molecular biology of the cell 25, 2677–2681 (2014).

5 Wani, M. C. & Horwitz, S. B. Nature as a Remarkable Chemist: A personal story of the discovery and development of Taxol®. Anti-cancer drugs 25, 482 (2014).

6 Suffness, M. & Wall, M. E. in Taxol 3–26 (CRC press, 2021).

7 Cech, N. B. & Oberlies, N. H. From plant to cancer drug: lessons learned from the discovery of taxol. Natural Product Reports 40, 1153–1157 (2023).

8 Wang, Y.-F. et al. Natural taxanes: developments since 1828. Chemical reviews 111, 7652–7709 (2011).

9 Shi, T. et al. An asymmetric catalytic multi-component reaction enabled the green synthesis of isoserine derivatives and semi-synthesis of paclitaxel. Green Synthesis and Catalysis 4, 58–63 (2023).

10 Min, L. et al. Strategies and Lessons Learned from Total Synthesis of Taxol. Chemical Reviews 123, 4934–4971 (2023).

11 Adhikari, P., Joshi, K. & Pandey, A. Taxus associated fungal endophytes: anticancerous to other biological activities. Fungal Biology Reviews 45, 100308 (2023).

12 Bureau, J. A., Oliva, M. E., Dong, Y. & Ignea, C. Engineering yeast for the production of plant terpenoids using synthetic biology approaches. Natural product reports 40, 1822–1848 (2023). 10.1039/d3np00005b

13 Iram, A., Dong, Y. & Ignea, C. Synthetic biology advances towards a bio-based society in the era of artificial intelligence. Curr Opin Biotechnol 87, 103143 (2024). 10.1016/j.copbio.2024.103143

14 Ajikumar, P. K. et al. Isoprenoid pathway optimization for Taxol precursor overproduction in Escherichia coli. Science 330, 70–74 (2010).

15 Malcı, K. et al. Improved production of Taxol® precursors in S. cerevisiae using combinatorial in silico design and metabolic engineering. Microbial Cell Factories 22, 243 (2023).

16 Nowrouzi, B., Torres-Montero, P., Kerkhoven, E. J., Martínez, J. L. & Rios-Solis, L. Rewiring Saccharomyces cerevisiae metabolism for optimised Taxol® precursors production. Metabolic Engineering Communications 18, e00229 (2024).

17 Jiang, B. et al. Characterization and heterologous reconstitution of Taxus biosynthetic enzymes leading to baccatin III. *Science*, eadj3484 (2024).

18 Zhang, Y. et al. Synthetic biology identifies the minimal gene set required for paclitaxel biosynthesis in a plant chassis. Molecular Plant 16, 1951–1961 (2023).

19 Lee, E.-K. et al. Cultured cambial meristematic cells as a source of plant natural products. Nature biotechnology 28, 1213–1217 (2010).

20 Hao da, C., Ge, G., Xiao, P., Zhang, Y. & Yang, L. The first insight into the tissue specific taxus transcriptome via Illumina second generation sequencing. PLoS One 6, e21220 (2011). 10.1371/journal.pone.0021220

21 Yu, C. et al. Identification of potential genes that contributed to the variation in the taxoid contents between two Taxus species (Taxus media and Taxus mairei). Tree Physiol 37, 1659–1671 (2017). 10.1093/treephys/tpx091

22 Li, S. T. et al. Transcriptional profile of Taxus chinensis cells in response to methyl jasmonate. BMC genomics 13, 295 (2012). 10.1186/1471-2164-13-295

23 Zhang, M. et al. Transcriptome-wide identification and screening of WRKY factors involved in the regulation of taxol biosynthesis in Taxus chinensis. Scientific reports 8, 5197 (2018). 10.1038/s41598-018-23558-1

24 Sun, G. et al. Deep sequencing reveals transcriptome re-programming of Taxus × media cells to the elicitation with methyl jasmonate. PLoS One 8, e62865 (2013). 10.1371/journal.pone.0062865

25 Jiao, Y. et al. Ancestral polyploidy in seed plants and angiosperms. Nature 473, 97–100 (2011). 10.1038/nature09916

26 Xiong, X. et al. The Taxus genome provides insights into paclitaxel biosynthesis. Nature plants 7, 1026–1036 (2021). 10.1038/s41477-021-00963-5

27 Leek, J. T. et al. Tackling the widespread and critical impact of batch effects in high-throughput data. Nature Reviews Genetics 11, 733–739 (2010).

28 Sprang, M., Andrade-Navarro, M. A. & Fontaine, J.-F. Batch effect detection and correction in RNA-seq data using machine-learning-based automated assessment of quality. BMC bioinformatics 23, 1–15 (2022).

29 Guo, F. et al. Concordance-Based Batch Effect Correction for Large-Scale Metabolomics. Analytical Chemistry 95, 7220–7228 (2023).

30 Bullard, J. H., Purdom, E., Hansen, K. D. & Dudoit, S. Evaluation of statistical methods for normalization and differential expression in mRNA-Seq experiments. BMC bioinformatics 11, 1–13 (2010).

31 Dillies, M.-A. et al. A comprehensive evaluation of normalization methods for Illumina high-throughput RNA sequencing data analysis. Briefings in bioinformatics 14, 671–683 (2013).

32 Abbas-Aghababazadeh, F., Li, Q. & Fridley, B. L. Comparison of normalization approaches for gene expression studies completed with high-throughput sequencing. PloS one 13, e0206312 (2018).

33 Vandenbon, A. Evaluation of critical data processing steps for reliable prediction of gene co-expression from large collections of RNA-seq data. Plos one 17, e0263344 (2022).

34 Thind, A. S. et al. Demystifying emerging bulk RNA-Seq applications: the application and utility of bioinformatic methodology. Briefings in bioinformatics 22, bbab259 (2021).

35 Conesa, A. et al. A survey of best practices for RNA-seq data analysis. Genome biology 17, 1–19 (2016).

36 Soneson, C., Love, M. I. & Robinson, M. D. Differential analyses for RNA-seq: transcript-level estimates improve gene-level inferences. F1000Research 4 (2015).

37 Sha, Y., Phan, J. H. & Wang, M. D. in 2015 37th Annual International Conference of the IEEE Engineering in Medicine and Biology Society (EMBC). 6461–6464 (IEEE).

38 Zhou, L., Sue, A. C.-H. & Goh, W. W. B. Examining the practical limits of batch effect-correction algorithms: When should you care about batch effects? Journal of genetics and genomics 46, 433–443 (2019).

39 Risso, D., Ngai, J., Speed, T. P. & Dudoit, S. Normalization of RNA-seq data using factor analysis of control genes or samples. Nature biotechnology 32, 896–902 (2014).

40 Goh, W. W. B., Wang, W. & Wong, L. Why batch effects matter in omics data, and how to avoid them. Trends in biotechnology 35, 498–507 (2017).

41 Goh, W. W. B., Yong, C. H. & Wong, L. Are batch effects still relevant in the age of big data? Trends in Biotechnology 40, 1029–1040 (2022).

42 Wang, Z., Gerstein, M. & Snyder, M. RNA-Seq: a revolutionary tool for transcriptomics. Nature reviews genetics 10, 57–63 (2009).

43 Evans, C., Hardin, J. & Stoebel, D. M. Selecting between-sample RNA-Seq normalization methods from the perspective of their assumptions. Briefings in bioinformatics 19, 776–792 (2018).

44 Bolger, A. M., Lohse, M. & Usadel, B. Trimmomatic: a flexible trimmer for Illumina sequence data. Bioinformatics 30, 2114–2120 (2014).

45 Bray, N. L., Pimentel, H., Melsted, P. & Pachter, L. Near-optimal probabilistic RNA-seq quantification. Nature biotechnology 34, 525–527 (2016).

46 Robinson, M. D. & Oshlack, A. A scaling normalization method for differential expression analysis of RNA-seq data. Genome biology 11, 1–9 (2010).

47 Robinson, M. D., McCarthy, D. J. & Smyth, G. K. edgeR: a Bioconductor package for differential expression analysis of digital gene expression data. bioinformatics 26, 139–140 (2010).

48 Law, C. W., Chen, Y., Shi, W. & Smyth, G. K. voom: Precision weights unlock linear model analysis tools for RNA-seq read counts. Genome biology 15, 1–17 (2014).

49 Love, M. I., Huber, W. & Anders, S. Moderated estimation of fold change and dispersion for RNA-seq data with DESeq2. Genome biology 15, 1–21 (2014).

50 Ritchie, M. E. et al. limma powers differential expression analyses for RNA-sequencing and microarray studies. Nucleic acids research 43, e47–e47 (2015).

51 Leek, J. T., Johnson, W. E., Parker, H. S., Jaffe, A. E. & Storey, J. D. The sva package for removing batch effects and other unwanted variation in high-throughput experiments. Bioinformatics 28, 882–883 (2012).

52 Smyth, G. K. in Bioinformatics and computational biology solutions using R and Bioconductor 397–420 (Springer, 2005).

53 Luo, J. et al. A comparison of batch effect removal methods for enhancement of prediction performance using MAQC-II microarray gene expression data. The pharmacogenomics journal 10, 278–291 (2010).

54 Dey, K. K., Hsiao, C. J. & Stephens, M. Visualizing the structure of RNA-seq expression data using grade of membership models. PLoS genetics 13, e1006599 (2017).

55 Levandowsky, M. & Winter, D. Distance between sets. Nature 234, 34–35 (1971).

56 Hubert, L. & Arabie, P. Comparing partitions. Journal of classification 2, 193–218 (1985).

57 Yadav, S. & Shukla, S. in 2016 IEEE 6th International conference on advanced computing (IACC). 78–83 (IEEE).

58 Pertea, G. & Pertea, M. GFF utilities: GffRead and GffCompare. F1000Research 9 (2020).

59 Conesa, A. et al. Blast2GO: a universal tool for annotation, visualization and analysis in functional genomics research. Bioinformatics 21, 3674–3676 (2005).

60 Xiong, X. et al. The Taxu s genome provides insights into paclitaxel biosynthesis. Nature Plants 7, 1026–1036 (2021).

61 Jiang, B. et al. Characterization and heterologous reconstitution of Taxus biosynthetic enzymes leading to baccatin III. Science 383, 622–629 (2024).

62 Abramson, J. et al. Accurate structure prediction of biomolecular interactions with AlphaFold 3. Nature 630, 493–500 (2024). 10.1038/s41586-024-07487-w

63 Kurre, D., Dang, P. X., Le, L. T. M., Gadkari, V. V. & Alam, A. Structural insights into binding-site access and ligand recognition by human ABCB1. The EMBO Journal 44, 991–1006 (2025). 10.1038/s44318-025-00361-z

64 Trott, O. & Olson, A. J. AutoDock Vina: improving the speed and accuracy of docking with a new scoring function, efficient optimization, and multithreading. J Comput Chem 31, 455–461 (2010). 10.1002/jcc.21334

65 Gutmann, D. A., Ward, A., Urbatsch, I. L., Chang, G. & van Veen, H. W. Understanding polyspecificity of multidrug ABC transporters: closing in on the gaps in ABCB1. Trends Biochem Sci 35, 36–42 (2010). 10.1016/j.tibs.2009.07.009

66 Janjic, A. et al. Prime-seq, efficient and powerful bulk RNA sequencing. Genome biology 23, 88 (2022).

67 Kim, S. et al. Genome sequence of the hot pepper provides insights into the evolution of pungency in Capsicum species. Nature genetics 46, 270–278 (2014).

68 Chen, W. et al. Whole-genome sequencing and analysis of the Chinese herbal plant Panax notoginseng. Molecular plant 10, 899–902 (2017).

69 Ignea, C. et al. Carnosic acid biosynthesis elucidated by a synthetic biology platform. Proc Natl Acad Sci U S A 113, 3681–3686 (2016). 10.1073/pnas.1523787113

70 Trikka, F. A. et al. Combined metabolome and transcriptome profiling provides new insights into diterpene biosynthesis in S. pomifera glandular trichomes. BMC genomics 16, 935 (2015). 10.1186/s12864-015-2147-3

71 Stegle, O., Parts, L., Piipari, M., Winn, J. & Durbin, R. Using probabilistic estimation of expression residuals (PEER) to obtain increased power and interpretability of gene expression analyses. Nature protocols 7, 500–507 (2012).

72 Kusano, H. et al. Taxus NPF transporter involved in the uptake of 10-deacetylbaccatin III facilitates the biosynthesis of taxane compounds. Plant J 122, e70146 (2025). 10.1111/tpj.70146

