## Supplementary Information for "Integrated RNA-seq analysis identifies ABC transporters mediating taxane export in *Taxus* species"

[#:] Contributed equally to this work

*Corresponding author:

### SUPPLEMENTARY FIGURES:

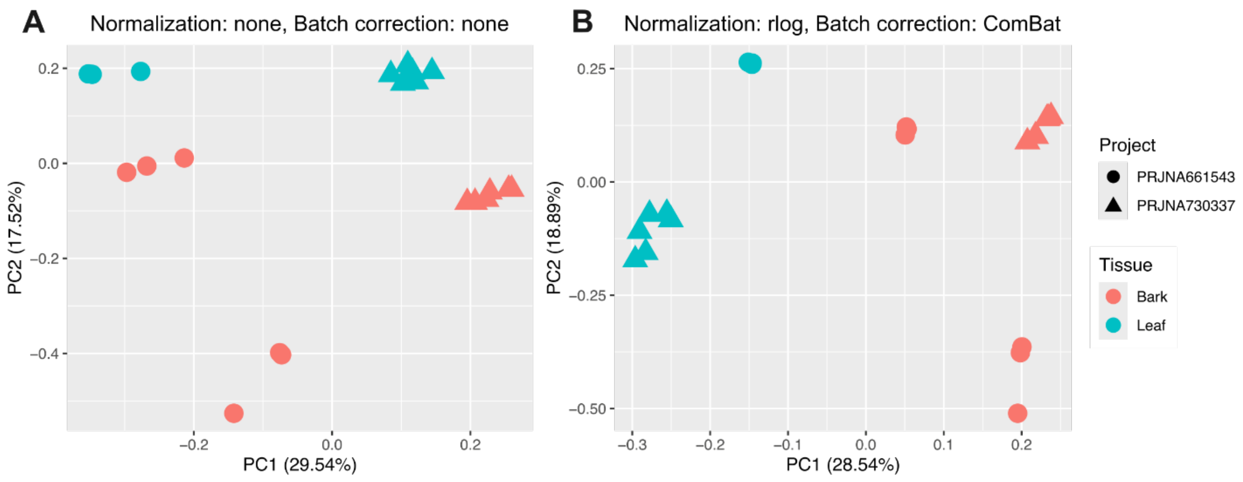


**Supplementary Figure. 1. PCA plots of expression data for the Bark+Leaf combined dataset A)** without normalization and batch effect correction, and **B**) after rlog normalization and ComBat BEC. n the unadjusted data (A), sample data cluster predominantly by project origin (**A**), indicating a strong influence of batch effect on the overall expression patterns. Upon application of rlog normalization and ComBat BEC, a stark shift in the clustering pattern occurs, with the now-corrected data aligning more closely by tissue type (**B**). This pronounced separation indicates that the combined normalization and BEC strategies can effectively mitigate the confounding batch variation and enhance the resolution of biologically relevant, tissue-specific expression patterns.

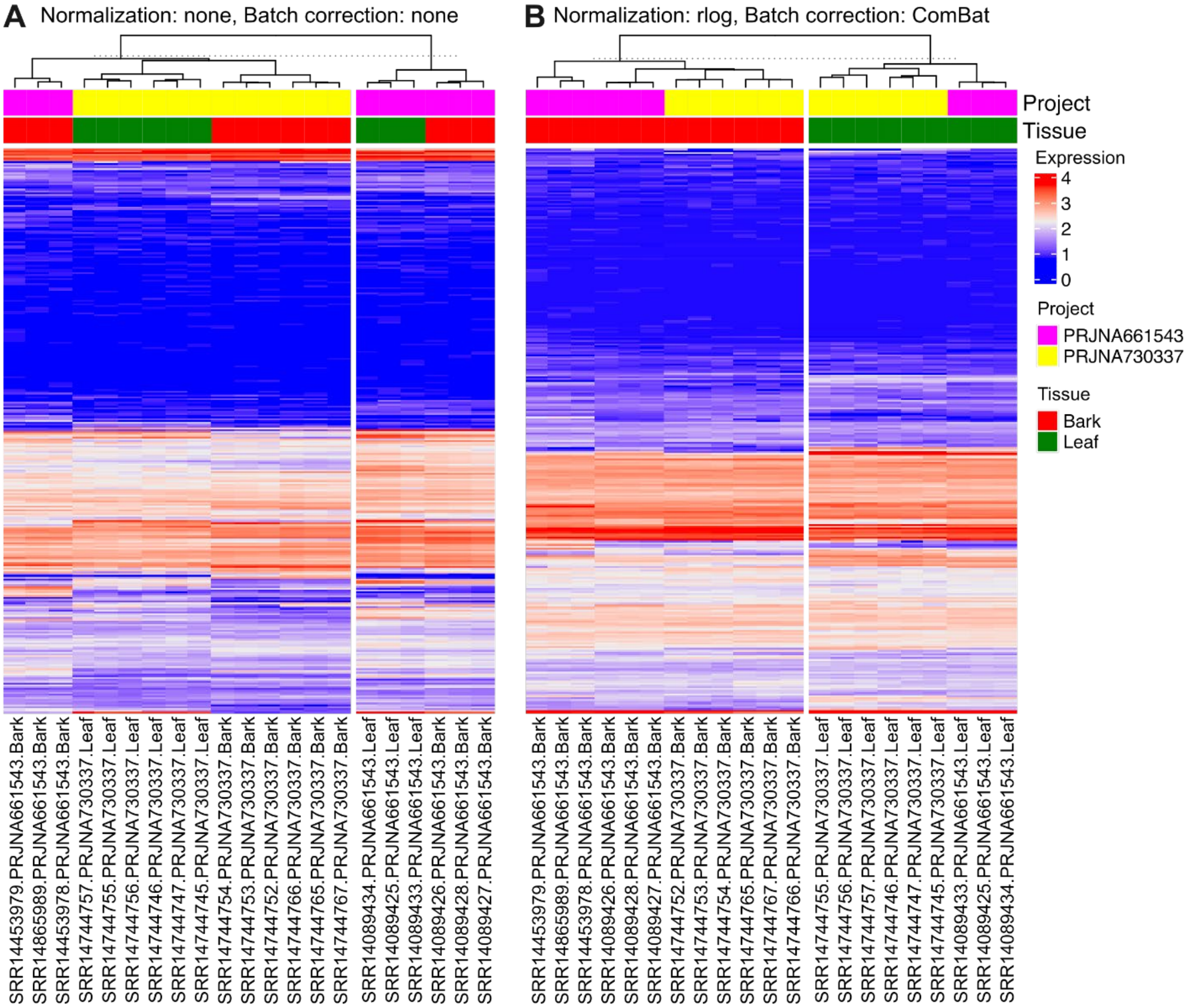


**Supplementary Figure 2. Representative clustering patterns illustrated by heatmap plots and *k*-means clustering** (k = 2) of the samples from the combined Bark+Leaf expression dataset **A**) without normalization and BEC, and **B**) after rlog normalization and ComBat batch effect correction. Prior to adjustment, the uncorrected data displayed a pronounced clustering by project, indicating a strong batch effect influencing the overall expression patterns (**A**). Following the application of rlog normalization and ComBat BEC, a striking shift in clustering was observed (**B**), with samples clearly segregated by tissue type.

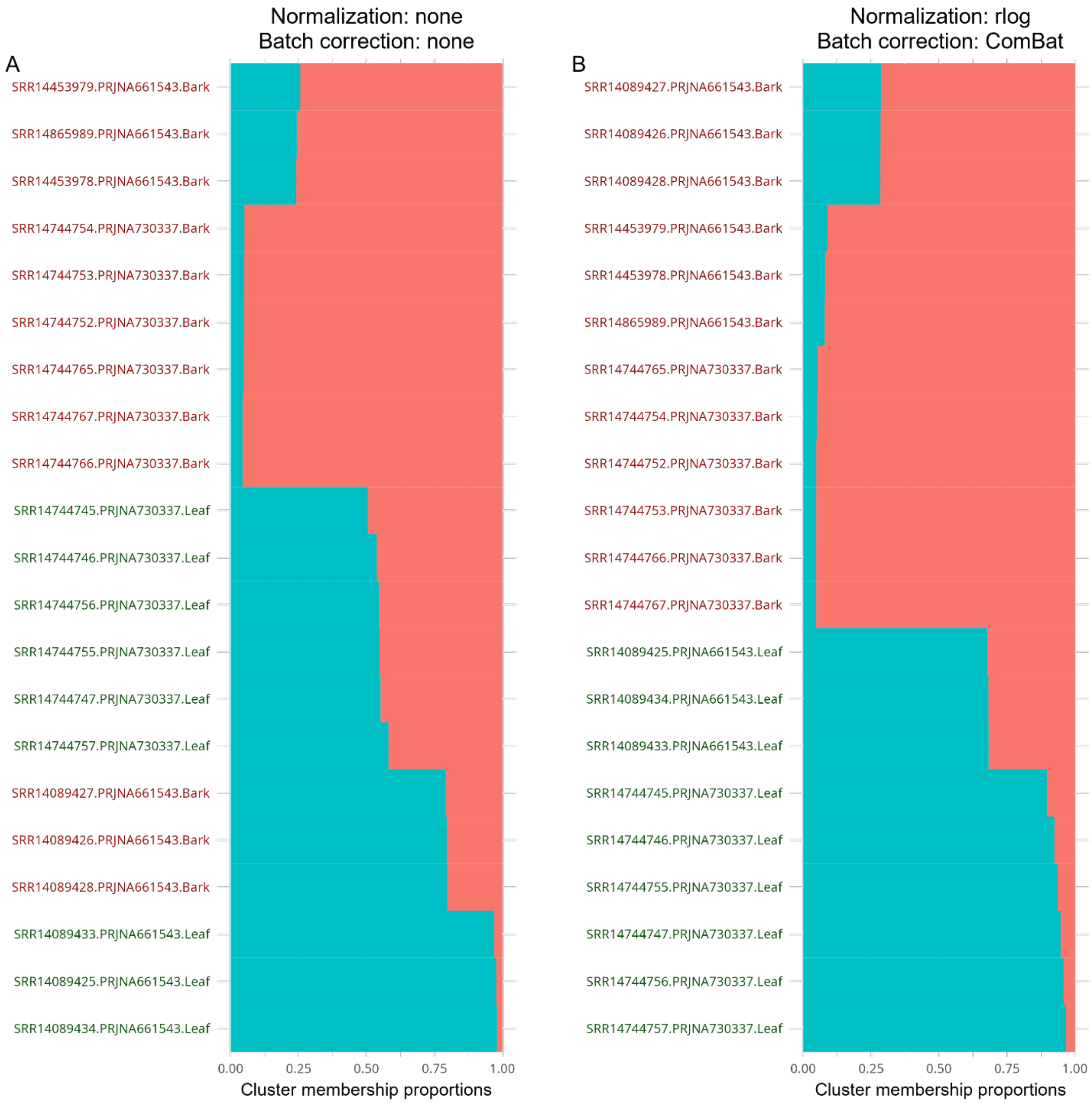


**Supplementary Figure 3.** Structure plots illustrating estimated membership proportions from a GoM model with k = 2 for the samples from the combined Bark+Leaf expression dataset: **A**) without normalization and BEC, and **B**) after rlog normalization and ComBat BEC. Each horizontal bar represents the cluster membership proportions of an individual sample. Prior to normalization and BEC, sample heterogeneity was evident, with no clear discrimination observed either for the project or for tissue based on cluster membership proportions. Following normalization and BEC, two distinct admixture patterns emerged, with blue indicating leaf samples and red indicating bark samples from two independent projects.

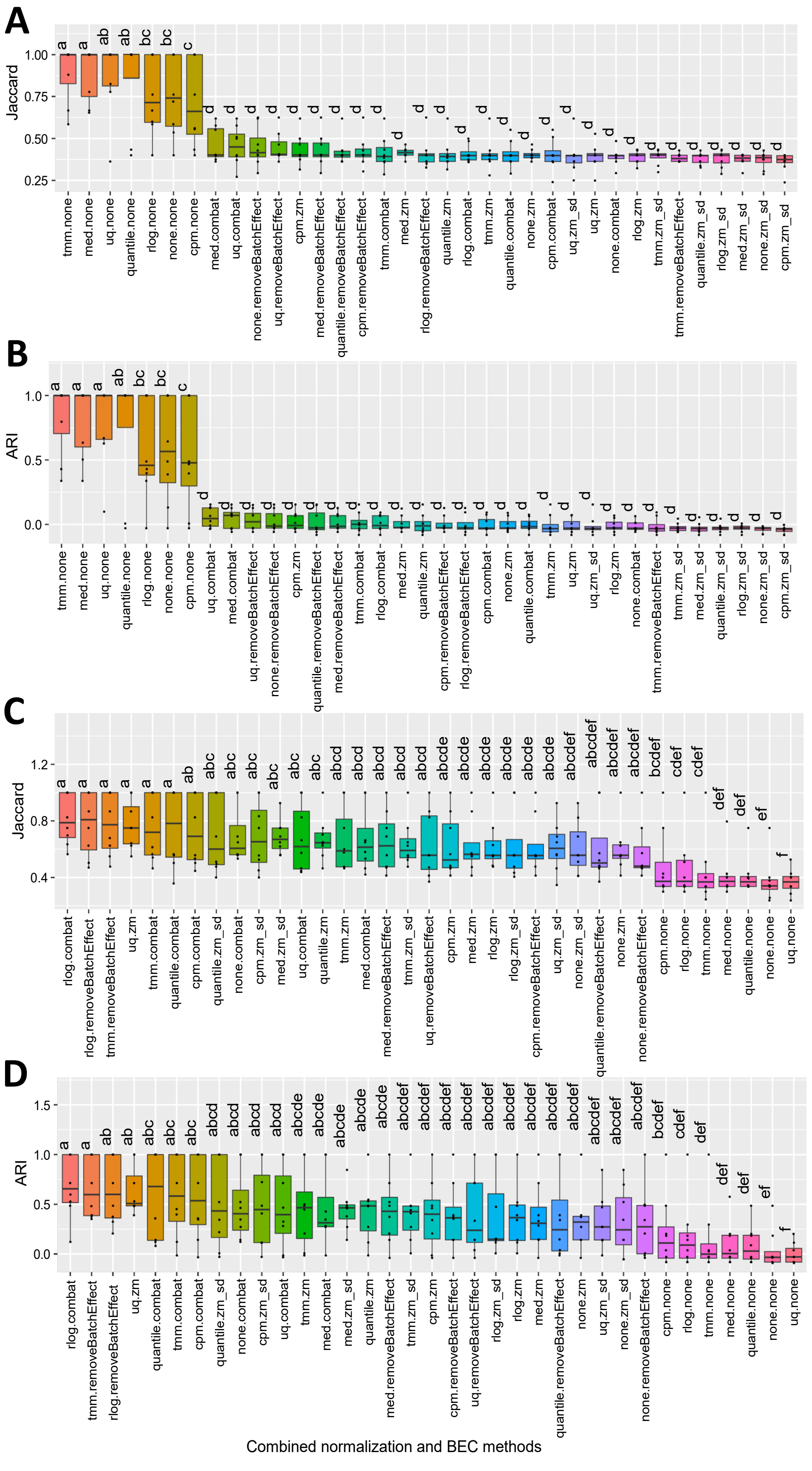
A
B
C
D
Jaccard
ARI
Combined normalization and BEC methods

**Supplementary Figure 4. Comparison of 35 different combinations using the box plot-assisted Duncan multiple range test for the *k*-means clustering method.** The combinations consist of 7 normalization methods crossed with 5 BEC approaches. Statistical significance was determined at $p < 0.05$ threshold, based on either the *Jaccard index* or *ARI*, with each box in the box plots containing 8 values. **A** and **B** represent the comparison of *k*-means cluster assignments to the project IDs (dataset of origin) assessed via the *Jaccard index* and *ARI*, respectively. **C** and **D** compare *k*-means cluster assignments to tissue types using the same metrics.

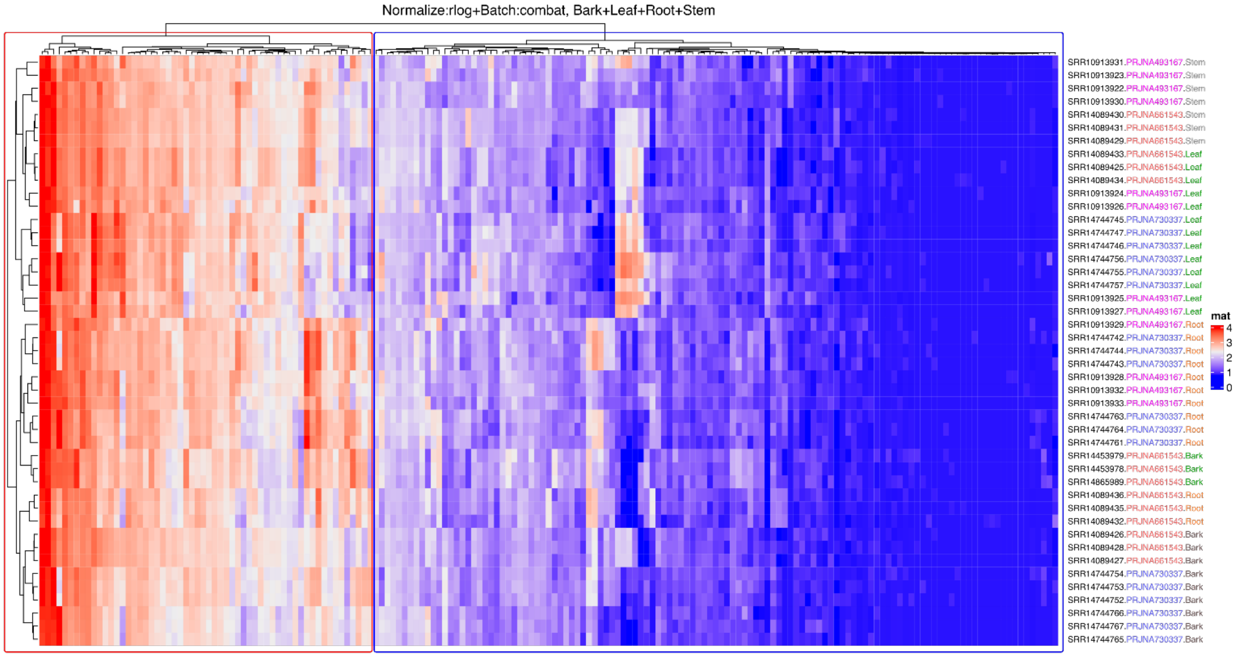


**Supplementary Figure 5. Clustered heatmap illustrating discrimination patterns of all putative ABC transporters across a combined tissue dataset** (bark, leaf, stem, and root). Notably, of the 189 putative ABC transporters, only 11 exhibited consistent expression levels, while the remaining 177 were grouped into a single cluster.

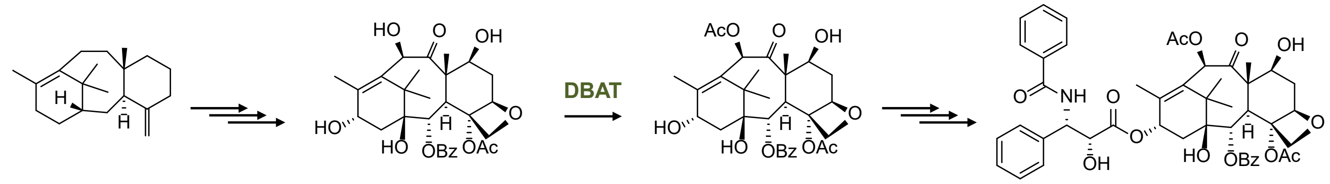


axa-4(20)-11(12)-diene | 10-deacetyl-baccatin III | baccatin III | taxol

**Supplementary Figure 6. Schematic of selective steps in the taxol biosynthetic pathway.** DBAT catalyzes the acetylation of 10-deacetyl-baccatin III, leading to the synthesis of the core taxol intermediate, baccatin III, the final intermediate before attachment of the functional side chain.
